# Predicting cell health phenotypes using image-based morphology profiling

**DOI:** 10.1101/2020.07.08.193938

**Authors:** Gregory P. Way, Maria Kost-Alimova, Tsukasa Shibue, William F. Harrington, Stanley Gill, Federica Piccioni, Tim Becker, Hamdah Shafqat-Abbasi, William C. Hahn, Anne E. Carpenter, Francisca Vazquez, Shantanu Singh

## Abstract

Genetic and chemical perturbations impact diverse cellular phenotypes, including multiple indicators of cell health. These readouts reveal toxicity and antitumorigenic effects relevant to drug discovery and personalized medicine. We developed two customized microscopy assays, one using four targeted reagents and the other three targeted reagents, to collectively measure 70 specific cell health phenotypes including proliferation, apoptosis, reactive oxygen species (ROS), DNA damage, and cell cycle stage. We then tested an approach to predict multiple cell health phenotypes using Cell Painting, an inexpensive and scalable image-based morphology assay. In matched CRISPR perturbations of three cancer cell lines, we collected both Cell Painting and cell health data. We found that simple machine learning algorithms can predict many cell health readouts directly from Cell Painting images, at less than half the cost. We hypothesized that these trained models can be applied to accurately predict cell health assay outcomes for any future or existing Cell Painting dataset. For Cell Painting images from a set of 1,500+ compound perturbations across multiple doses, we validated predictions by orthogonal assay readouts, and by confirming mitotic arrest, ROS, and DNA damage phenotypes via PLK, proteasome, and aurora kinase/tubulin inhibition, respectively. We provide an intuitive web app to browse all predictions at http://broad.io/cell-health-app. Our approach can be used to add cell health annotations to Cell Painting perturbation datasets.

## Introduction

Perturbing cells with specific genetic and chemical reagents in different environmental contexts impacts cells in various ways (Kitano, 2002). For example, certain perturbations impact cell health by stalling cells in specific cell cycle stages, increasing or decreasing proliferation rate, or inducing cell death via specific pathways (Markowetz, 2010; Szalai et al., 2019). Cell health is normally assessed by eye or measured by specifically targeted reagents, which are either focused on a single Cell Health parameter (ATP assays) or multiple, in combination, via FACS-based or image-based analyses, which involves a manual gating approach, complicated staining procedures, and significant reagent cost. These traditional approaches limit the ability to scale to large perturbation libraries such as candidate compounds in academic and pharmaceutical screening centers.

Image-based profiling assays are increasingly being used to quantitatively study the morphological impact of chemical and genetic perturbations in various cell contexts (Caicedo et al., 2016; Scheeder et al., 2018). One unbiased assay, called Cell Painting, stains for various cellular compartments and organelles using non-specific and inexpensive reagents (Gustafsdottir et al., 2013). Cell Painting has been used to identify small-molecule mechanisms of action (MOA), study the impact of overexpressing cancer mutations, and discover new bioactive mechanisms, among many other applications (Caicedo et al., 2018; Christoforow et al., 2019; Hughes et al., 2020; Pahl and Sievers, 2019; Rohban et al., 2017; Simm et al., 2018; Wawer et al., 2014). Additionally, Cell Painting can predict overall mammalian toxicity levels for environmental chemicals (Nyffeler et al., 2020) and some of its derived morphology measurements are readily interpreted by cell biologists and relate to cell health (Bray et al., 2016). However, no single, inexpensive assay enables discovery of fine-grained cell health readouts that would provide researchers with a more complete understanding of perturbation mechanisms.

We hypothesized that we could predict many cell health readouts directly from the Cell Painting data, which is already available for hundreds of thousands of perturbations. This would enable the rapid and interpretable annotation of small molecules or genetic perturbations. To do this, we first developed two customized microscopy assays, which collectively report on 70 different cell health indicators via a total of seven reagents applied in two reagent panels. Collectively, we call these assays “Cell Health”.

To demonstrate proof of concept, we collected a small pilot dataset of 119 CRISPR knockout perturbations in three different cell lines using Cell Painting and Cell Health. We used the Cell Painting morphology readouts to train 70 different regression models to predict each Cell Health indicator independently. We used simple machine learning methods instead of a deep learning approach because of our limited sample size of 119 perturbations and the inability to increase the sample size by linking single cell measurements across assays. We predicted certain readouts, such as the *number of S phase cells*, with high performance, while performance on other readouts, such as *DNA damage in G2 phase cells*, was low. We applied and validated these models on a separate set of existing Cell Painting images acquired from 1,571 compound perturbations measured across six different doses from the Drug Repurposing Hub project (Corsello et al., 2017). We provide all predictions in an intuitive web-based application at http://broad.io/cell-health-app, so that others can extend our work and explore cell health impacts of specific compounds.

## Results and Discussion

We collected Cell Painting images and targeted Cell Health readouts in three different cell lines (A549, ES2, HCC44), each treated with 119 CRISPR perturbations targeting 59 genes and controls (**Supplementary Table 1**). We selected these genes to span multiple biological pathways and induce different morphological states. The seven reagents we included in the two Cell Health panels **(Supplementary Table 2**) include specific stains and antibodies such as Caspase 3/7 dye to target apoptotic cells and γH2Ax antibodies to measure DNA damage.

Applying biological knowledge of cell-health related phenotypes and several manual gating strategies, we defined 70 different Cell Health readouts (**Supplementary Table 3**) based on signals from the seven reagents, plus nucleus morphology measurements from Digital Phase Contrast (DPC) (**Figure 1A**, **Supplementary Figure 1, Supplementary Figure 2**). While these readouts are relatively easy to interpret, running two separate assays is not ideal for large-scale perturbation screening experiments.

**Figure 1.**
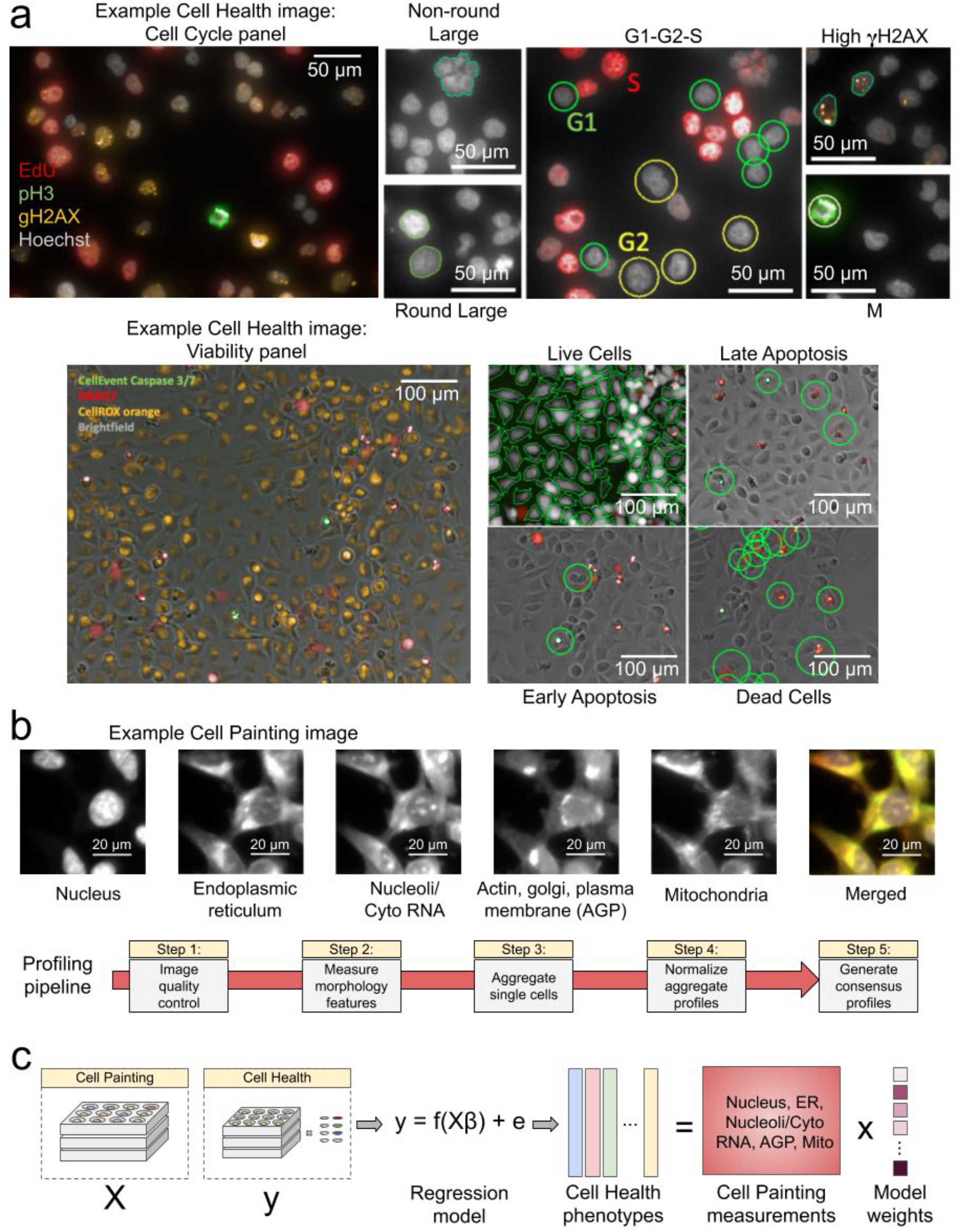
Data processing and modeling approach. **(a)** Example images and workflow from the Cell Health assays. We apply a series of manual gating strategies (see Methods) to isolate cell subpopulations and to generate cell health readouts for each perturbation. (top) In the “Cell Cycle” panel, in each nucleus we measure Hoechst, EdU, PH3, and gH2AX. (bottom) In the “Cell Viability” panel, we capture digital phase contrast images, measure Caspase 3/7, DRAQ7, and CellROX. **(b)** Example Cell Painting image across five channels, plus a merged representation across channels. The image is cropped from a larger image and shows ES2 cells. Scale bars are 20 μm. Below are the steps applied in an image-based profiling pipeline, after features have been extracted from each cell’s image. **(c)** Modeling approach where we fit 70 different regression models using CellProfiler features derived from Cell Painting images to predict Cell Health readouts. Model weights refer to the coefficients derived from each regression model.

We developed and applied a bioinformatics pipeline to process features extracted from Cell Painting images by CellProfiler software (Carpenter et al., 2006). The pipeline yields image-based profiles representing gene and guide perturbation signatures (Caicedo et al., 2017) (**Figure 1B**). We observed that 63% of guide profile replicates were distinguishable from negative controls; that is, they had stronger pairwise correlations than 95% of a null distribution defined by non-replicate correlations (**Supplementary Figure 3**). This rate is consistent with previous Cell Painting studies of genetic perturbations (Rohban et al., 2017).

We developed an approach to use the inexpensive reagents from the multiplexed, high-throughput Cell Painting assay to predict Cell Health readouts (**Figure 1C**). We generated a single, “consensus” signature for each guide perturbation across cell lines, producing 357 signatures (3 cell lines x 119 CRISPR guides) with 952 morphology measurements. We independently optimized 70 different elastic net linear regression models using consensus morphology profiles of Cell Painting data to predict each of the 70 Cell Health readouts independently (**Figure 1C**). The actual identity of the CRISPR guides were not relevant during training.

Predictive performance in a held-out test set (a balanced 15% of profiles not used in training) indicates high expected generalizability for many models (**Figure 2, Supplementary Figure 4**). Performance was better for nearly every model when trained with real data compared to shuffled data, thus beating a random chance baseline (**Supplementary Figure 5**).

**Figure 2:**
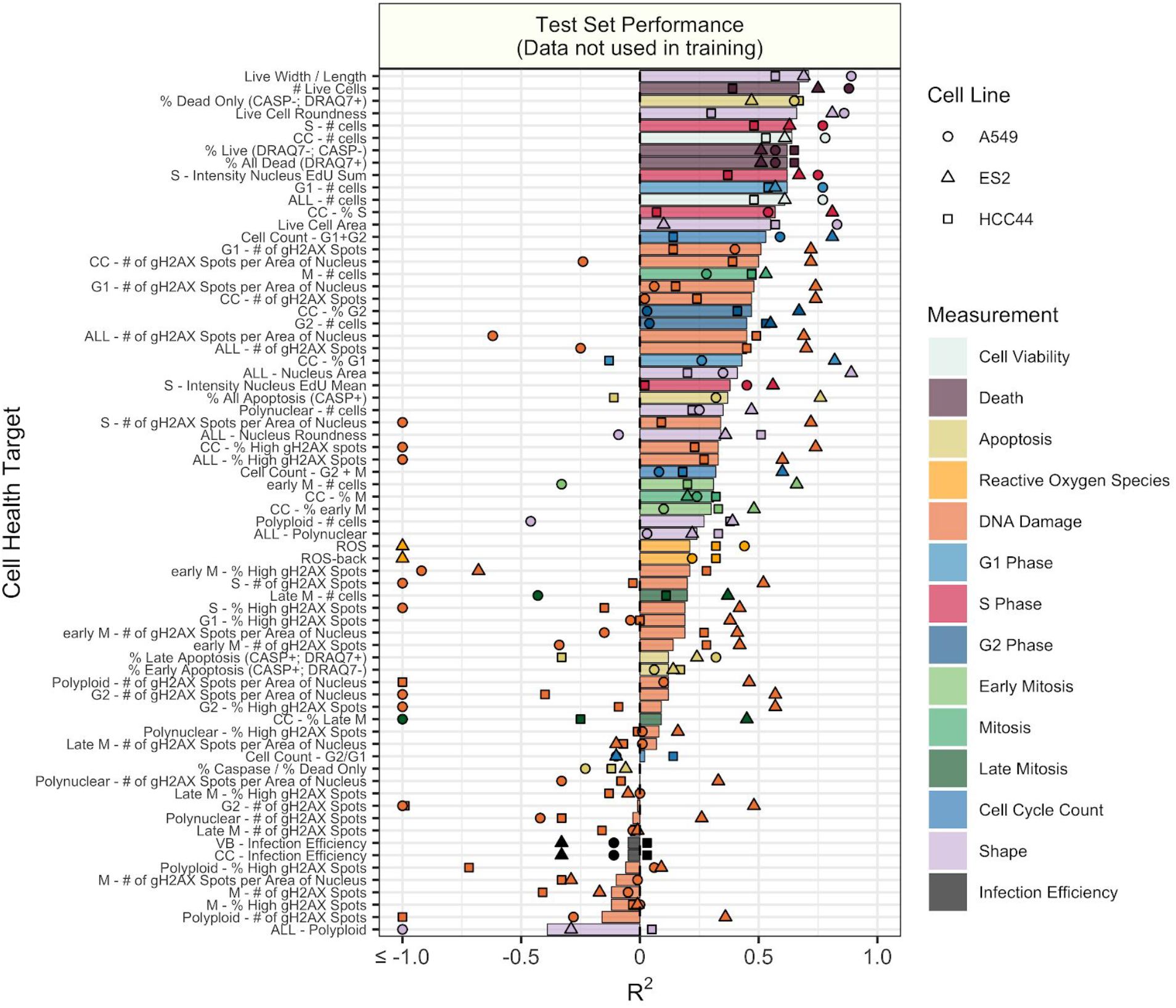
Test set model performance of predicting 70 Cell Health readouts with independent regression models. Performance for each phenotype is shown, sorted by decreasing R^2^ performance. The bars are colored based on the primary measurement metadata (see **Supplementary Table S3**), and they represent performance aggregated across the three cell lines. The points represent cell line specific performance. Points falling below −1 are truncated to −1 on the x axis. See **Supplementary Figure 3** for a full depiction.

Many Cell Health readouts were predicted very well, including *percentage of dead cells, number of S-phase cells*, *DNA damage in G1-phase cells*, and *percentage of apoptotic cells* (**Supplementary Figure 6A**). However, other readouts such as *DNA damage in polynuclear cells*, and *percentage of cells in late mitosis* could not be predicted better than random (**Supplementary Figure 6B**). Models derived from different combinations of Cell Health reagents had variable performance, with DRAQ7, shape, and EdU models performing the best (**Supplementary Figure 7**). Performance differences might result from random technical variation, small sample sizes for training models, different number of cells in certain Cell Health subpopulations (e.g. mitosis or polynuclear cells), fewer cells collected in the viability panel (see methods), or the inability of Cell Painting reagents to capture certain phenotypes. We observed overall better predictivity in ES2 cells, which had the highest CRISPR infection efficiency (**Supplementary Figure 8**), suggesting that stronger perturbations provide better information for training and that training on additional data should provide further benefit.

Furthermore, using a linear model for predictions enables interpretability. For example, inspecting the model for the Cell Health readout *Live Cell Area* reveals that it relies on cell and cytoplasm shape features from Cell Painting (**Supplementary Figure 9**). This is expected given that the *Live Cell Area* readout is derived from cell boundary measurements from the digital phase contrast channel. In our approach, each regression model uses a combination of interpretable morphology features to make Cell Health phenotype predictions, unlike so-called “black box” deep learning feature extractors. Therefore, the specific combination of Cell Painting features provides a potentially interpretable morphology signature representing the underlying cell health state.

Overall, many different feature classes were important for accurate predictions (**Figure 3, Supplementary Figure 10**). Some features tended to strongly contribute across multiple Cell Health readouts. For example, particularly informative features include the radial distribution of the actin, golgi, and plasma membrane (AGP) channel in cells and DNA granularity in nuclei. This demonstrates that the Cell Painting assay captures complex cell health phenotypes using a rich variety of morphology feature types.

**Figure 3:**
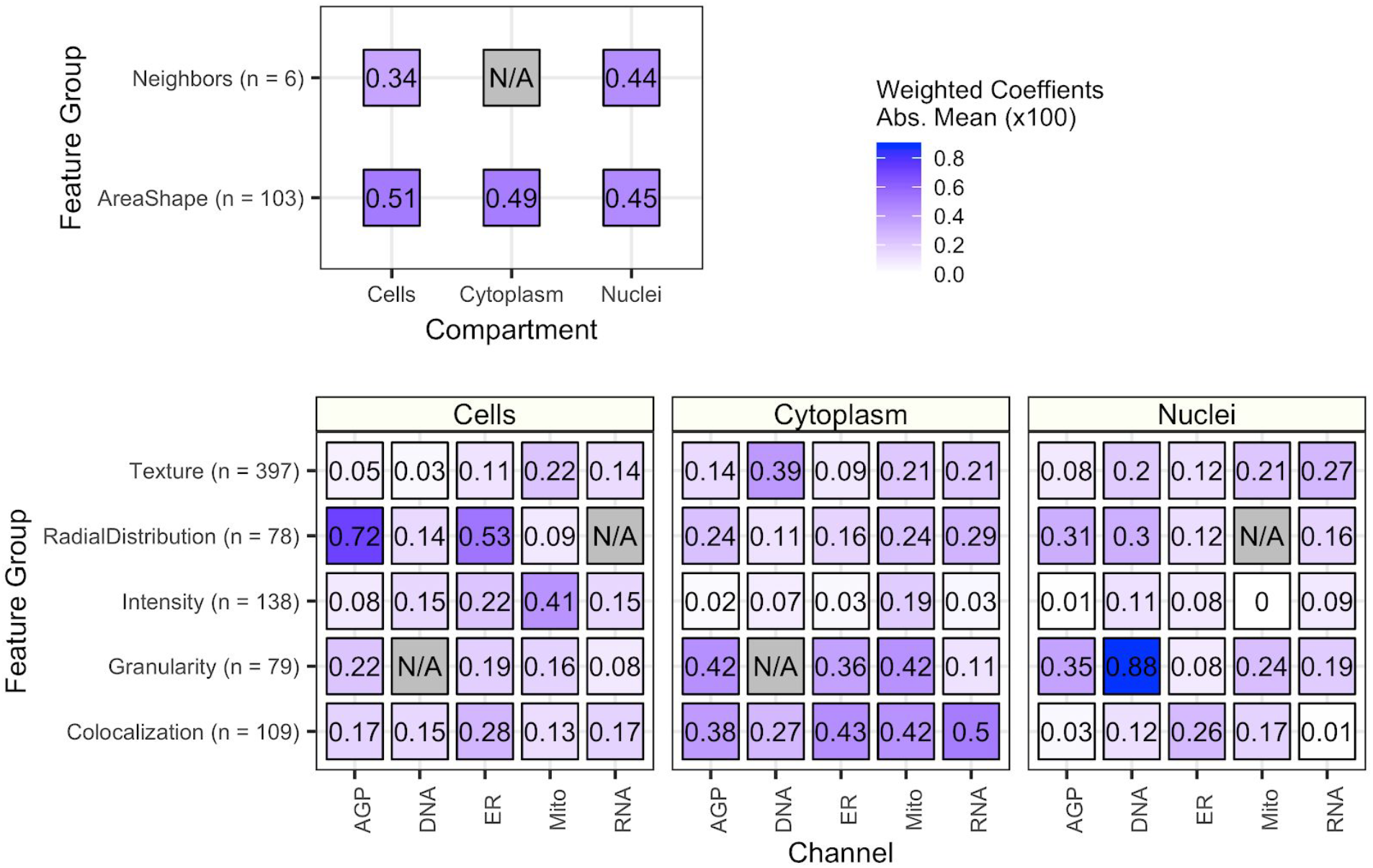
The importance of each class of Cell Painting features in predicting 70 Cell Health readouts. Each square represents the mean absolute value of model coefficients weighted by test set R^2^ across every model. The features are broken down by compartment (Cells, Cytoplasm, and Nuclei), channel (AGP, Nucleus, ER, Mito, Nucleolus/Cyto RNA), and feature group (AreaShape, Neighbors, Channel Colocalization, Texture, Radial Distribution, Intensity, and Granularity). The number of features in each group, across all channels, is indicated. For a complete description of all features, see the handbook: http://cellprofiler-manual.s3.amazonaws.com/CellProfiler-3.0.0/index.html. Dark gray squares indicate “not applicable”, meaning either that there are no features in the class or the features did not survive an initial preprocessing step. Note that for improved visualization we multiplied the actual model coefficient value by 100.

We performed a series of analyses to determine certain parameters and options that are likely to improve models in the future. First, we performed a “cell line holdout” analysis, in which we trained models on two of three cell lines and predicted cell health readouts on the held out cell line. We observed that certain models including those based on viability, S phase, early mitotic and death phenotypes could be moderately predicted in cell lines agnostic to training (**Supplementary Figure 11**). Not surprisingly, shape-based phenotypes could not be predicted in holdout cell lines, which emphasizes the limitations of transferring certain cell-line intrinsic measurements across cell lines. We also performed a systematic feature removal analysis, in which we retrained cell health models after dropping features that are measured from specific groups, compartments, and channels. We observed that many models were robust to dropping entire feature classes during training (**Supplementary Figure 12**). This result demonstrates that many Cell Painting features are highly correlated, which might permit prediction “rescue” even if the directly implicated morphology features are not measured. Because of this, we urge caution when generating hypotheses regarding causal relationships between phenotypes and individual Cell Painting features. Lastly, we performed a sample size titration analysis in which we randomly removed a decreasing amount of samples from training. For the high and mid performing models we observed a consistent performance drop, suggesting that increasing sample size would result in better overall performance (**Supplementary Figure 13**).

Predictive models of cell health would be most useful if they could be trained once and successfully applied to data sets collected separately from the experiment used for training. Otherwise one could not annotate existing datasets that lack parallel Cell Health results, and Cell Health assays would have to be run alongside each new dataset. We therefore applied our trained models to a large, publicly-available Cell Painting dataset collected as part of the Drug Repurposing Hub project (Corsello et al., 2017). The data derive from A549 lung cancer cells treated with 1,571 compound perturbations measured in six doses.

We first chose a high-performing model to validate. The *number of live cells* model captures the number of cells that are unstained by DRAQ7. We compared model predictions to orthogonal viability readouts from a third dataset: Publicly available PRISM assay readouts, which count barcoded cells after an incubation period (Yu et al., 2016). Despite measuring perturbations with slightly different doses and being fundamentally different ways to count live cells (**Figure 4A**), the predictions correlated with the assay readout (Spearman’s Rho = 0.35, p < 1 x 10^−3^; **Figure 4B**).

**Figure 4:**
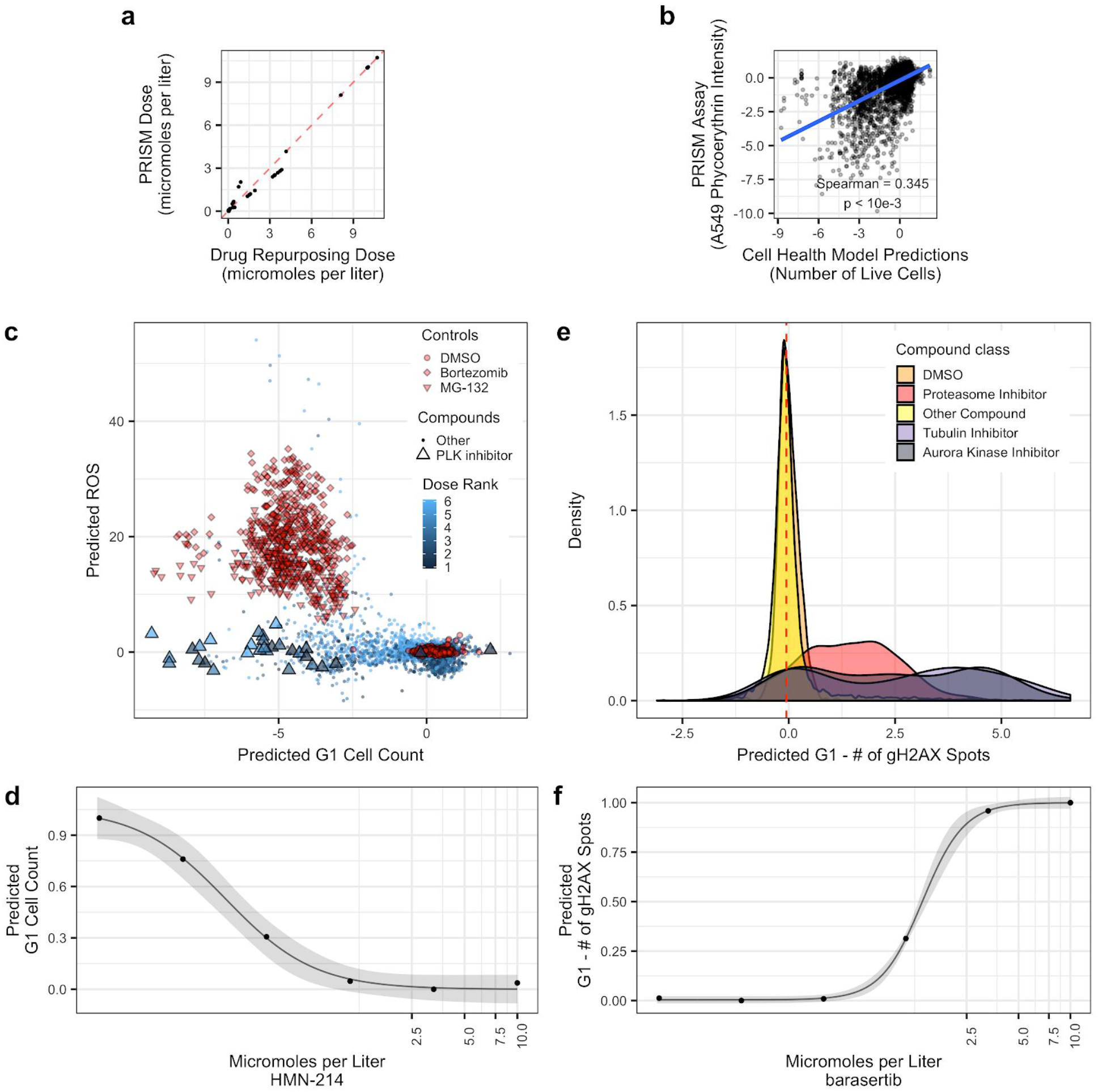
Validating Cell Health models to Cell Painting data from The Drug Repurposing Hub. The models were not trained using the Drug Repurposing Hub data. **(a)** The results of the dose alignment between the PRISM assay and the Drug Repurposing Hub data. This view indicates that there was not a one-to-one matching between perturbation doses. **(b)** Comparing viability estimates from the PRISM assay to the predicted number of live cells in the Drug Repurposing Hub. The PRISM assay estimates viability by measuring barcoded A549 cells after an incubation period. **(c)** Drug Repurposing Hub profiles stratified by *G1 cell count* and *ROS* predictions. Bortezomib and MG-132 are proteasome inhibitors and are used as positive controls in the Drug Repurposing Hub set; DMSO is a negative control. We also highlight all PLK inhibitors in the dataset. **(d)** HMN-214 is an example of a PLK inhibitor that shows strong dose response for *G1 cell count* predictions. **(e)** Tubulin and aurora kinase inhibitors are predicted to have high Number of gH2AX spots in G1 cells compared to other compounds and controls. **(f)** Barasertib (AZD1152) is an aurora kinase inhibitor that is predicted to have a strong dose response for Number of gH2AX spots in G1 cells predictions.

We also chose to validate three additional models: *ROS, G1 cell count*, and *Number of gH2AX spots in G1 cells*. We observed that the two proteasome inhibitors (bortezomib and MG-132) in the Drug Repurposing Hub set yielded high *ROS* predictions (OR = 76.7; p < 1 x 10^−15^) (**Figure 4C**) Proteasome inhibitors are known to induce ROS (Han and Park, 2010; Ling et al., 2003). As well, PLK inhibitors yielded low *G1 cell counts* (OR = 0.035; p = 3.9 x 10^−8^) (**Figure 4C**). The PLK inhibitor HM-214 showed an appropriate dose response (**Figure 4D**). PLK inhibitors block mitotic progression, thus reducing entry into the G1 cell cycle phase (Lee et al., 2014). Lastly, we observed that aurora kinase and tubulin inhibitors yielded high *Number of gH2AX spots in G1 cells* predictions (OR = 11.3; p < 1 x 10-15) (**Figure 4E**). In particular, we observed a strong dose response for the aurora kinase inhibitor barasertib (AZD1152) (**Figure 4F**). Aurora kinase and tubulin inhibitors cause prolonged mitotic arrest, which can lead to mitotic slippage, G1 arrest, DNA damage, and senescence (Cheng and Crasta, 2017; Orth et al., 2011; Tsuda et al., 2017).

We applied uniform manifold approximation (UMAP) to observe the underlying structure of the samples as captured by morphology data (McInnes et al., 2018). We observed that the UMAP space captures gradients in predicted *G1 cell count* (**Supplementary Figure S14A**) and in predicted *ROS* (**Supplementary Figure S14B**). We also observed similar gradients in the ground truth cell health readouts in the CRISPR Cell Painting profiles used for training cell health models (**Supplementary Figure S15**). Gradients in our data suggest that cell health phenotypes manifest in a continuum rather than in discrete states.

Lastly, we observed moderate technical artifacts in the Drug Repurposing Hub profiles, indicated by high DMSO profile dispersion in the Cell Painting UMAP space (**Supplementary Figure 14C**); this represents an opportunity to improve model predictions with new batch effect correction tools. Additionally, it is important to note that the expected performance of each Cell Health model can only be as good as the performance observed in the original test set (see **Figure 2**), and that all predictions require further experimental validation.

## Conclusions

We have demonstrated feasibility that information in Cell Painting images can predict many different Cell Health indicators even when trained on a relatively small dataset. The results motivate collecting larger datasets for training, with more perturbations and multiple cell lines. These new datasets would enable the development of more expressive models, based on deep learning, that can be applied to single cells. Including orthogonal imaging markers of CRISPR infection would also enable us to isolate cells with expected morphologies. More data and better models would improve the performance and generalizability of Cell Health models and enable annotation of new and existing large-scale Cell Painting datasets with important mechanisms of cell health and toxicity.

## Methods

### CRISPR constructs used for knockout

We performed a clustered regularly interspersed short palindromic repeats (CRISPR) and CRISPR-associated protein 9 (Cas9) knockout experiment to perturb cells (CRISPR-Cas9). We designed guides to target 59 different genes with an average of two guides per gene (**Supplementary Table 1**). All sgRNAs were selected from the Avana library (Doench et al., 2016; Meyers et al., 2017) or by using CRISPick (https://broad.io/crispick) (Hanna and Doench, 2020). Nevertheless, it is important to note that the identity of the CRISPR gene target is not used in training the machine learning models.

### Cell lines

We performed CRISPR knockout in three different cell lines (A549, ES2, and HCC44). All cell lines used were stably expressing Cas9 and were part of the Achilles project (Meyers et al 2017). Prior to data collection, we confirmed cell line identity using single nucleotide polymorphism (SNP) profiling. We confirmed that all cell lines were mycoplasma negative by using MycoAlert Mycoplasma detection kits (Lonza, Walkersville, MD).

### Lentiviral infection and plating

Virus was prepared in 96-well plate according to the published protocol (https://portals.broadinstitute.org/gpp/public/resources/protocols). Before initiating the screen we optimized the number of cells per well and polybrene concentration for each Cas9-expressing cell line. Ultimately, we plated A549, ES2, and HCC44 cells with starting densities of 350, 375, and 150 cells per well, respectively in 384-well black-wall, clear-bottom plates (Corning Costar). We also optimized sgRNA lentivirus volume to achieve 100% infection while maintaining low toxicity. For the screen, we spin-infected cells with 4 ug/ML polybrene concentration at the optimized density and virus volume (Aguirre et al., 2016). Three parallel plates were seeded per cell line. On one plate, cells were treated with or without 2 ug/ml puromycin 24 hours post-infection, and cell viability was determined using CellTiterGlo (Promega) after 96 hours of selection to determine infection efficiency. The second and third plates were used for the Cell Health assays (cell cycle and viability).

### Cell Painting: Cell staining and image acquisition

We followed the traditional Cell Painting protocol to acquire the readouts (Bray et al., 2016). We treated the cells with CRISPR guide perturbations and incubated for five days. Following the incubation period, we fixed cells with 10 μl of 16% (wt/vol) methanol-free paraformaldehyde (PFA) for a final concentration of 3.2% (vol/vol). We imaged cells with a PerkinElmer Opera Phenix confocal HCI microscope at 20x magnification. We applied the standard panel of Cell Painting dyes to mark various cellular compartments: Hoechst 33342 to mark DNA, Concanavalin A/Alexa 488 to mark endoplasmic reticulum (ER), SYTO 14 to mark the nucleoli and cytoplasmic RNA, Phalloidin/Alexa 568 and wheat-germ agglutinin/Alexa 555 to mark actin cytoskeleton, golgi, and plasma membrane (AGP), and MitoTracker Deep Red/Alexa 647 to mark mitochondria (Bray et al., 2016). We collected nine images per well in five different channels for these different unbiased stains. In total, we collected 138,226 pictures after quality control filtering, which includes five channels per site, nine sites per well, across nine 384-well plates. In total, this represents about 2 TB of images. We deposited raw and illumination corrected images to the Image Data Resource (https://idr.openmicroscopy.org) under accession number idr0080 (Williams et al., 2017).

### Cell Painting: Image processing

The next step in a Cell Painting protocol is to extract morphology features from the images that can be used as an unbiased systems biology measurement to describe how each perturbation impacts various cellular compartments in the assay. We built a CellProfiler image analysis and illumination correction pipeline (version 2.2.0) pipeline to extract these image-based features (McQuin et al., 2018). We include the CellProfiler pipelines in our github repository. Using the CellProfiler pipeline, we first performed several adjustments to account for potential confounding factors such as background intensity and illumination correction. Next, we used our pipeline to segment cells, distinguish between nuclei and cytoplasm, and then measure specific features related to the various channels captured. We measured the fluorescence intensity, texture, granularity, density, location, and various other measurements for each single cell (see http://cellprofiler-manual.s3.amazonaws.com/CellProfiler-3.0.0/index.html for more details). Following the image-analysis pipeline, we obtain 8,964,210 cells and 1,785 feature measurements across 9 different plates. We provide the raw output single cell profiles as extracted from our CellProfiler pipeline on figshare (Way et al., 2019).

### Cell Painting: Image-based profiling

After the image analysis pipeline, the next step is to process the single cell image-based features that are output of the CellProfiler pipeline. We used a standard approach (Caicedo et al., 2017) to process the single cell profiles. First, we aggregated all single cells grouped by perturbation (effectively, by well) by computing the median value per morphology feature. This process takes all single cells and computes a single perturbation profile that is used to compare all perturbations against each other downstream. Next, using the median and median absolute deviation of feature values from empty wells as the center and scale parameters respectively, we normalized all perturbation profiles by subtracting the center and dividing by the scale, and did so for each plate independently. This normalization procedure transforms all features to exist on the same scale and enables the perturbation profiles to be compared across plates and batches.

We then applied a feature selection procedure to reduce noisy and retain the most informative features. We removed features with missing values in any profile, features with low variance, features with extreme outlier values, and blocklisted features. Extreme outlier features are defined by having measurements greater than 15 standard deviations following normalization. The blocklisted features are generally unreliable features that are known to be noisy and have caused numerical issues in previous experiments (Way, 2020). We used pycytominer (https://github.com/cytomining/pycytominer) to perform the profiling pipeline, which can be reproduced at https://github.com/broadinstitute/cell-health.

Following these procedures we derived profiles for 357 perturbations representing 119 guides measured across the three different cell lines. We also computed the perturbation consensus signatures of the Cell Painting data (see Methods: Forming consensus signatures). Our final Cell Painting dataset had 357 consensus profiles measured by 952 morphology features (357 x 952). These data are available on github https://github.com/broadinstitute/cell-health/tree/master/1.generate-profiles/data/profiles.

### Cell Health assay: Cell staining and image acquisition

We treated all cells with a panel of specific reagents each measuring a different aspect of cell health (see **Supplementary Table S2**). The seven reagents include unbiased dyes, click chemistry, and specific antibody treatments. The reagents measure various aspects of cell health including proliferation, mitosis, DNA damage, reactive oxygen species (ROS), and apoptosis timing. We collected a minimum of four replicates per treatment. Because many reagents fluoresce in different emission spectra, we applied the reagents in parallel. We applied a series of semi-manual gating strategies to isolate specific cell health phenotypes in specific cell subsets (**Supplementary Table S3**). Together, we refer to the collection of measurements as the “Cell Health” assays.

More specifically, we collect the Cell Health assay data in a series of two distinct panels: Cell cycle and viability. In the first panel we measure Hoechst, EdU, PH3, and gH2AX, and use these measurements to quantify cell cycle and DNA damage in specific cell cycle subsets. In the second panel, we measure viability phenotypes using Caspase 3/7, DRAQ7, CellRox, and digital phase contrast (DPC) nucleus morphology measurements.

We acquired all cell images using an Opera Phenix HCI Instrument (PerkinElmer) with a 20X water objective (a numerical aperture (NA) of 1.0), in confocal mode. We acquired images in four channels using default excitation / emission combinations: for the blue channel (Hoechst) 405/435-480; for the green channel (Alexa 488 and CellEvent) 488/500-550; for the orange channel (Alexa 568 and CellRox Orange) 561/570-630 and for the far-red channel (Alexa 647 and DRAQ7) 640/650-760. We applied the Cell Health reagents for cell viability and for cell cycle in two separate plates.

The first set of plates (n = 3 replicate plates) measures cell cycle. We added 5-ethynyl-2’-deoxyuridine (EdU) in live cells for S phase cells to integrate. We then fixed the cells with 4% formaldehyde using standard approaches and detected EdU using Click-iT™ EdU Alexa Fluor™ 647 HCS Assay (Thermo Fisher C10357) according to the vendor protocol. We then performed standard immunofluorescence (IF) staining with two antibodies: one targeting phosphohistone H3 (PH3) to measure cells undergoing mitosis and one to identify DNA damage foci in nuclei via γH2Ax. We followed these PH3 and γH2Ax antibody treatments by secondary antibodies conjugated with Alexa 488 and Alexa 568, respectively. We added Hoecsht 33342 dye to stain nuclear DNA. For the cell cycle plate, we acquired nine fields of view per well.

The second set of plates (n = 3 replicate plates) measures cell viability. We added CellEvent™ Caspase-3/7 Green Detection Reagent (ThermoFisher), DRAQ7, and CellROX™ Orange Reagent (ThermoFisher) dyes to measure apoptotic cells, dead cells, and reactive oxygen species (ROS), respectively. We acquired one field of view per well using green, orange and far-red fluorescence channels as well as brightfield and Digital Phase Contrast (DPC) channels. The cells were incubated at 37C.

### Cell Health assay: Image analysis

We developed and ran two distinct image analysis pipelines in Harmony software (version 4.1; PerkinElmer) for each of the Cell Health plates. Individually for each cell line assayed, and for both cell cycle and cell viability plates, we established a series of manual gating strategies to identify distinct cell line subpopulations (see **Figure 1A**).

For the cell cycle plate, we performed nucleus segmentation using the Hoechst channel and discarded all nuclei that were at the field border. We identified cells in specific cell cycle stages using Hoechst, Alexa 488 (pH3) and Alexa 647 (EdU) intensities. We identified γH2AX spots within nuclei based on the Alexa 568 channel. These strategies are standard in the field (Aguirre et al., 2016). More specifically, we identified subpopulations based on the respective channel intensities and morphological properties for each nucleus as specified:

- We stratified populations “polyploid”, “polynuclear (large not round nuclei)”, and “cells selected for cell cycle” based on total intensity of the Hoechst channel (DNA content) and nucleus “roundness” measurements as output from the PerkinElmer Harmony software.
- We identified four subpopulations within the “cells selected for cell cycle” population as follows:

- “G1 Cells”: selected based on low total Hoechst intensity and low green (pH3) and far red (EdU) channels. We excluded outlier nuclei with unusually high intensity of Hoechst max.
- “G2 Cells”: Selected based on high total Hoechst intensity and low green (pH3) and far red (EdU) channels. We excluded outlier nuclei with unusually high intensity of Hoechst max.
- “G2/M Cells”: Selected based on the same criteria as for G2, except we included nuclei with high green (pH3) mean.
- “M Cells”: Selected based on high green (pH3) mean.
- “S Cells”: Selected based on high mean far red (EdU) channel intensity.
- We counted orange spots (γH2AX) representing DNA damage loci in each of the cell cycle subpopulations. We determined high γH2AX activity if there were more than three spots per nucleus.

For the cell viability plate, we performed cell segmentation based on the cumulative Digital Phase Contrast (DPC) channels. Again, we identified specific subpopulations based on the following channel intensities:

- We separated “Dead Cells” and “Live Cells” based on max intensity far red channel (DRAQ7).

- We identified a “Dead Only Cells” subpopulation within the “Dead Cells” population by isolating cells without green (Caspase 3/7) signal.
- We identified “Caspase Positive Cells” based on green (Caspase 3/7) channel max intensity.

- We distinguished two subpopulations in the “Caspase Positive Cells” named “Early Apoptotic Cells” and “Late Apoptotic Cells” based on low and high far red (DRAQ7) max signal intensity, respectively.
- We used the mean intensity of the CellROX™ Orange signal to measure ROS.

- We excluded edge wells in the ROS analysis because of consistent poor signal quality.

Additionally, we set these gates for each cell subpopulation using a set of random wells from each cell line and experiment independently. We observed that the intensity measurements used to form the gates were consistent across wells and plates, and generally formed distinct cell subpopulation clusters. After using the random wells to set the gates, we used the Harmony microscope software to apply the gates to the remaining wells and plates.

We also used CRISPR infection efficiency, which is measured in a separate assay for both cell cycle and viability imaging assays, as negative control features. In total, considering both plates and all cell subpopulations, we measured 70 different variables in the Cell Health Assay. To standardize plate-level differences, we normalized Cell Health readouts per plate by subtracting median values and dividing by the standard deviation.

### Forming consensus signatures

After acquiring the images and processing the data, we prepared the data further before input into our machine learning framework. We generated consensus signatures for each perturbation using a moderated z-score (MODZ) procedure (Subramanian et al., 2017). Briefly, we calculated pairwise Spearman correlations between all replicates of a single perturbation and then combined profiles by weighting their signature contribution by mean pairwise correlation to all other replicates. We applied this transformation to both Cell Painting profiles and Cell Health assay measurements. We collected more replicates of the Cell Painting data than the Cell Health data. In total, we collected assay readouts from 357 common perturbations (119 CRISPR guides across three cell lines). In the Cell Painting data, we filtered and collapsed 3,456 morphology profiles to the common set of 357 consensus profiles. In the Cell Health assays, we filtered and collapsed 2,302 well profile readouts to the common set of 357 consensus readouts. We generated consensus signatures because there is no way to match replicate-level information across the assays. We applied UMAP (McInnes et al., 2018) to the consensus profiles and visualized patterns of ground truth cell health measurements.

### Machine learning framework

We randomly split 15% of the consensus signatures into a separate test set. We balanced this stratification by cell line. We then used the remaining 85% to train all 70 cell health models. In total, we used 303 samples as the training set and 54 as the test set.

We elected to train elastic net regression models using *sklearn* (version 0.20.3) (Pedregosa et al., 2011; Zou and Hastie, 2005). We chose this model because it is quick to train, is easily interpretable, and will induce sparsity in selecting model features. We also trained classification models and binarized training and testing data by using >1.5 standard deviations away from the mean as positive examples. However, because the classification approach was unstable and sensitive to low sample sizes, we elected to move forward using only the regression models (see https://github.com/broadinstitute/cell-health/issues/78). To identify optimal alpha and elastic net mixing parameters, we performed a grid search and 5-fold cross validation using the training set only. For each model independently, we observed cross validation performance across 9 different alpha parameters ([0.01, 0.1, 0.2, 0.3, 0.4, 0.5, 0.6, 0.7, 0.8]) and 11 different elastic net mixing parameters ([0.1, 0.12, 0.14, 0.16, 0.2, 0.3, 0.4, 0.5, 0.7, 0.8, 0.9]). Alpha controls the regularization penalty term for all features, and the elastic net mixing parameter controls the trade-off between L1 and L2 regression where 0 = L1 and 1 = L2. Therefore, the closer the elastic net mixing parameter is to 0, the sparser the model. We optimized and trained 70 different elastic net regression models for each of the 70 Cell Health assay readouts independently.

We repeated this procedure and independently optimized 70 additional models using randomly shuffled data. For the shuffling procedure, we randomly shuffled the Cell Painting features independently per column before training. We use the shuffled model performance as a suitable baseline to compare real model performance.

### Machine learning evaluation

We evaluated each of the 70 Cell Health regression models independently using R squared statistics from *sklearn* version 0.20.3 (Pedregosa et al., 2011). We calculated R squared for the full training and testing partitions, in shuffled training and testing partitions, and for each cell line independently for all 70 models. The measurement can be interpreted as how well the models could predict the real Cell Health readout with values approaching one as perfect fits. It is best to compare test set performance in real vs. shuffled data. The test set performance in real data simulates how models are expected to perform in data not used for model training. The shuffled performance indicates if there is any expected performance inflation.

### Machine learning robustness: Investigating the impact of sample size

We performed an analysis in which we randomly dropped an increasing amount of samples from the training set before model training. After dropping the predefined number of samples, we retrained all 70 cell health models and assessed performance on the original holdout test set. We performed this procedure ten times with ten unique random seeds to mirror a more realistic scenario of new data collection and to reduce the impact of outlier samples on model training.

### Machine learning robustness: Systematically removing feature classes

We performed an analysis in which we systematically dropped features measured in specific compartments (Nuclei, Cells, and Cytoplasm), specific channels (RNA, Mito, ER, DNA, AGP), and specific feature groups (Texture, Radial Distribution, Neighbors, Intensity, Granularity, Correlation, Area Shape) and retrained all models. We omitted one feature class and then independently optimized all 70 cell health models as described in the Machine learning framework results section above. We repeated this procedure once per feature class.

### Drug Repurposing Hub Cell Painting data: Imaging-based profiling

A subset of the Drug Repurposing Hub compounds (n = 1,571) (Corsello et al., 2017) were profiled using the Cell Painting assay across about six doses per compound. We processed this dataset using a standard image-based profiling pipeline to extract consensus profiles per treatment. See https://github.com/broadinstitute/lincs-cell-painting for complete details and instructions on how to reproduce. Briefly, we applied a CellProfiler image analysis pipeline to segment cells, adjust for background intensity, and measure morphology features for three compartments: cells, cytoplasm, and nuclei. The output of this procedure was 136 SQLite files (one for each plate) representing unnormalized single-cell profiles. Next, we developed and applied an image-based profiling bioinformatics pipeline to generate treatment consensus profiles from the single-cell measurements (Caicedo et al., 2017). The same image analysis pipeline and bioinformatics pipeline were used to process all the plates in the experiment.

In the pipeline, we first median-aggregated the single cells by feature to form well profiles, and then, using the median and median absolute deviation of feature values from dimethyl sulfoxide (DMSO) as the center and scale parameters respectively, we normalized all perturbation profiles by subtracting the center and dividing by the scale, and did so for each plate independently.

The plates in this dataset have 24 DMSO-treated wells and therefore represented a good alignment control to adjust for plate level differences. Each plate also has two positive controls (BRD-K50691590 (Bortezomib) and BRD-K60230970 (MG-132)) at 20 millimoles per liter with 12 replicates each for all plates. We visualized positive and negative control profiles in our UMAP space to determine the extent of technical artifacts present in our data. Following the z-score normalization, we combined all treatment replicates (~6 per compound and dose pair) using MODZ consensus signatures. We generated consensus profiles for control replicates by well, across plate maps. In total, this procedure resulted in 10,752 different treatment profiles and 1,788 normalized CellProfiler morphology features.

### Drug Repurposing Hub Cell Painting data: Predicting Cell Health readouts

We applied all Cell Health models to the 10,752 Drug Repurposing Hub consensus Cell Painting profiles. We simply applied the Cell Health trained models using the *sklearn model.predict()* method. Every feature measured in the CRISPR perturbation Cell Painting profiles were also measured in the Drug Repurposing Hub output. The result of the model application was 70 Cell Health readouts for all 10,752 treatments. We used these predictions for model validation with external data, and for visualization in the web app scatter plots.

### Assessing generalizability of cell health models applied to Drug Repurposing Hub data

We used our cell health webapp (https://broad.io/cell-health-app) to identify compounds with high predictions for three models with high or intermediate performance: *ROS, Number of G1 cells*, and *Number of gH2AX spots in G1 cells*. For each model, we identified classes of compounds with consistently high scores, then tested for statistical enrichment: for proteasome inhibitors in the *ROS* model, PLK inhibitors in the *Number of G1 cells* model, and aurora kinase and tubulin inhibitors in the *Number of gH2AX spots in G1 cells* model. We used one-sided Fisher’s exact tests to quantify differences in expected proportions between high and low model predictions. For each case, we determined high and low predictions based on the 50% quantile threshold for each model independently.

### Drug Repurposing Hub Cell Painting data: Visualization

We also applied UMAP to the 10,752 Drug Repurposing Hub Cell Painting profiles and extracted two lower dimensional representations. UMAP reduces the Cell Painting profiles to two features that capture the global structure of the input data. Prior to UMAP transformation, we applied a feature selection procedure to the Drug Repurposing Hub profiles. We removed features with low variance, features with missing values in any consensus profile, blocklisted features (Way, 2020), and features with extreme outlier values defined by greater than 15 standard deviations following normalization. This procedure reduced the feature dimension from 1,788 to 572. By reducing the number of features, we can be more confident that the major sources of variation are not biased by CellProfiler feature redundancies or by technical artifacts of sample processing.

### Drug Repurposing Hub Cell Painting data: Dose-response analysis

To model dose, we fit Hill equations (4 parameter log-logistic model) to all 1,571 Drug Repurposing compounds consensus signatures transformed into each of the 70 different Cell Health model predictions. Before input into the model, we zero-one transformed Cell Health predictions across doses for each compound independently. For most compounds, this normalization procedure happens for six data points (representing six doses per compound consensus signature) at a time. The zero-one procedure assigns a value of zero to the lowest value, one to the highest value, and scales each intermediate value accordingly. This procedure results in 109,970 different model fits. We used the *drc* R package (version 3.0-1) to fit all models (Ritz et al., 2015). We present all precomputed dose fit models to be explored in the broad.io/cell-health-app.

### Comparing viability predictions to an orthogonal readout

We downloaded the PRISM assay results (version 19Q4) from the Cancer Dependency Map website at https://depmap.org/portal/download/ (Corsello et al., 2017). The PRISM assay measures viability of multiple cell lines in a pooled format and deconvolutes results based on barcoded readouts (Yu et al., 2016). We focused on the A549 cell line and compounds that were measured in both the PRISM assay and Drug Repurposing collection. The PRISM assay profiled 1,382 of the 1,571 Drug Repurposing Compounds. The PRISM assay also used slightly different doses than the Drug Repurposing Hub collection procedure. Therefore, to align doses, we converted doses into dose ranks, and report Spearman correlations between the two datasets (see **Figure 4A**).

### Code and data availability

All data and code are publicly available. Analysis software to reproduce the full paper is available at https://github.com/broadinstitute/cell-health/tree/v2.0 (Way et al., 2020). Raw and illumination corrected Cell Painting images are available in the Image Data Resource (accession number idr0080). Single cell morphology profiles derived from these images are available at NIH Figshare at https://doi.org/10.35092/yhjc.9995672.v5. Processed Cell Painting profiles, and raw and processed Cell Health readouts are also available at https://github.com/broadinstitute/cell-health/tree/v2.0. Processing code and data for the Cell Painting Drug Repurposing Hub data is available at https://github.com/broadinstitute/lincs-cell-painting/tree/v0.1. Cell Health predictions for the Drug Repurposing Hub compounds are available to explore at https://broad.io/cell-health-app.

## Supporting information

Supplementary Table S1 - Perturbation Details

Supplementary Table S2 - Cell Health Assay Reagents

Supplementary Table S3 - Annotated Cell Health Measurements

## Acknowledgements

We would like to thank Kyle Karhohs for executing the image analysis pipeline, Nasim Jamali, David Stirling, and Beth Cimini for insightful discussions about CellProfiler features and cell images, Kate Hartland for inputs on staining protocols, and the LINCS data generators in the Broad Institute’s Connectivity Map Team for producing the Drug Repurposing Hub Cell Painting data.

## Supplementary Figures

**Supplementary Figure S1.**
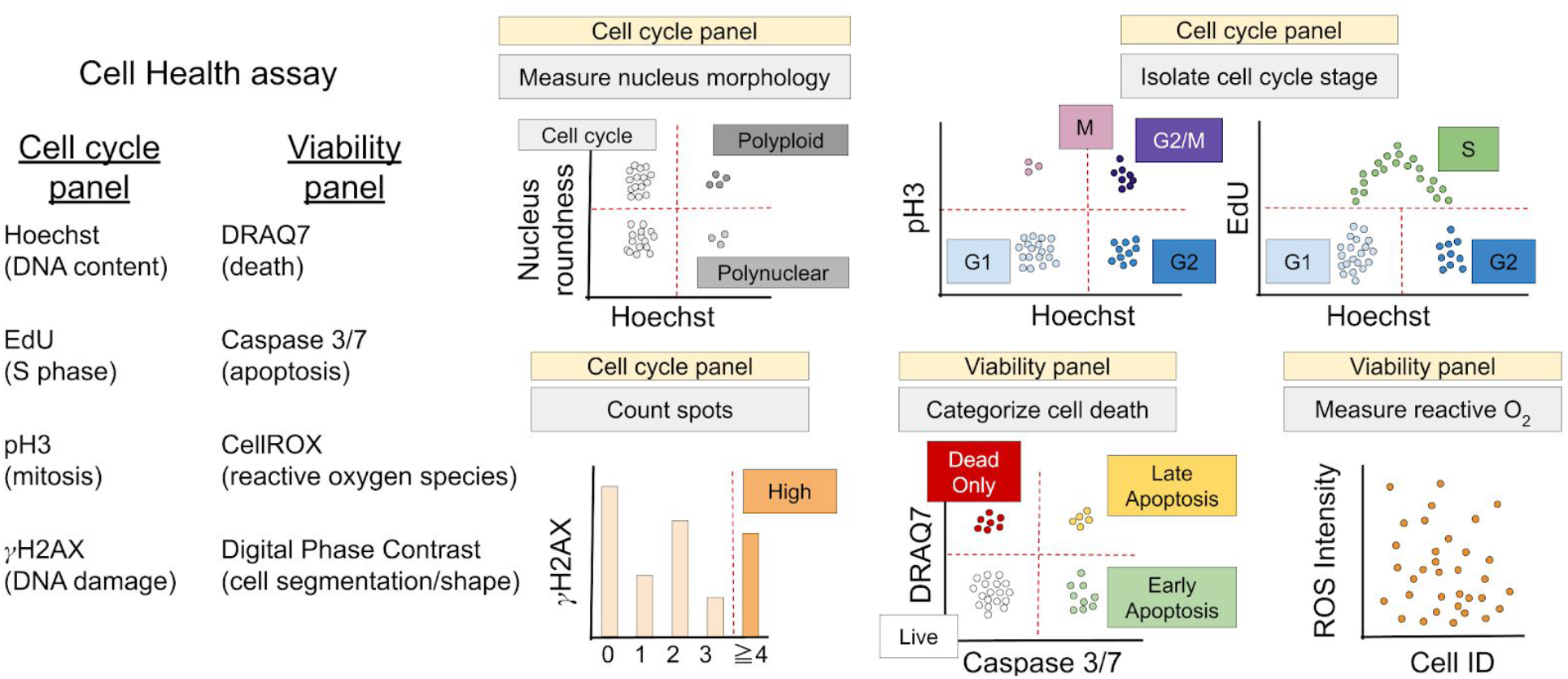
Illustration of the gating strategy in the Cell Health assays. We extract 70 different readouts from the Cell Health imaging assays. The assay consists of two customized reagent panels, which use measurements from seven different targeted reagents and one channel based on digital phase contrast (DPC) imaging; shown are five toy examples to demonstrate that individual cells are isolated into subpopulations by various gating strategies to define the Cell Health readouts.

**Supplementary Figure S2.**
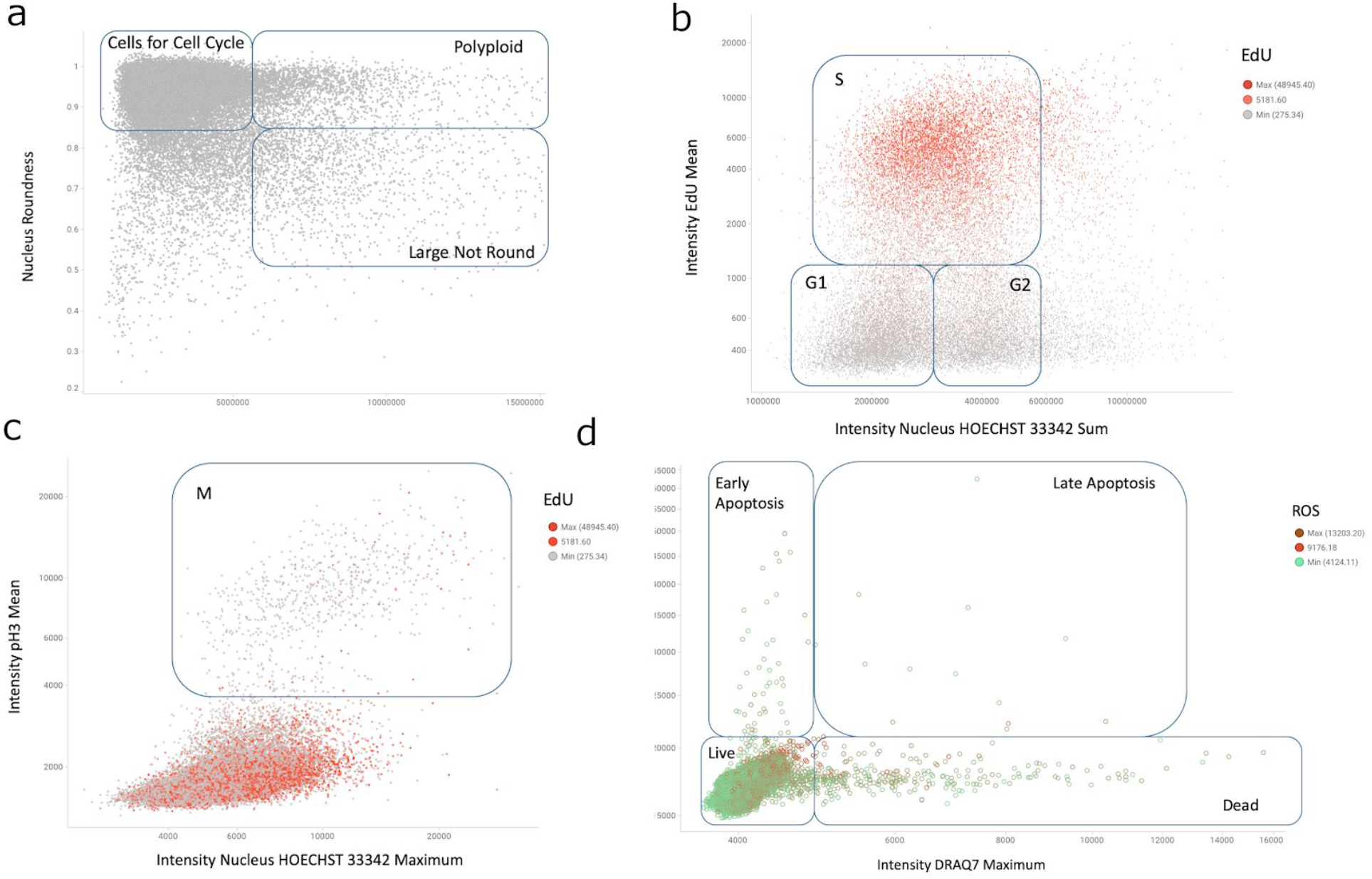
Real data of manual gating in the Cell Health assays. For each cell line, we apply a series of manual gating strategies defined by various stain measurements in single cells to define cell subpopulations. **(a)** In the cell cycle panel, we first select cells that are useful for cell cycle analysis based on nucleus roundness and Hoechst intensity measurements. We also identify polyploid and “large not round” (polynuclear) cells. **(b)** We then subdivide the cells used for cell cycle to G1, G2, and S cells based on total Hoechst intensity (DNA content) and EdU incorporation signal intensity. **(c)** We use Hoechst and PH3 nucleus intensity to define mitotic cells. The points are colored by EdU intensity in the nucleus in both (b) and (c). **(d)** Example gating in the viability panel. We use DRAQ7 and CellEvent (Caspase 3/7) to distinguish alive and dead cells, and categorize early or late apoptosis. See Methods for more details about how the Cell Health measurements are made.

**Supplementary Figure S3.**
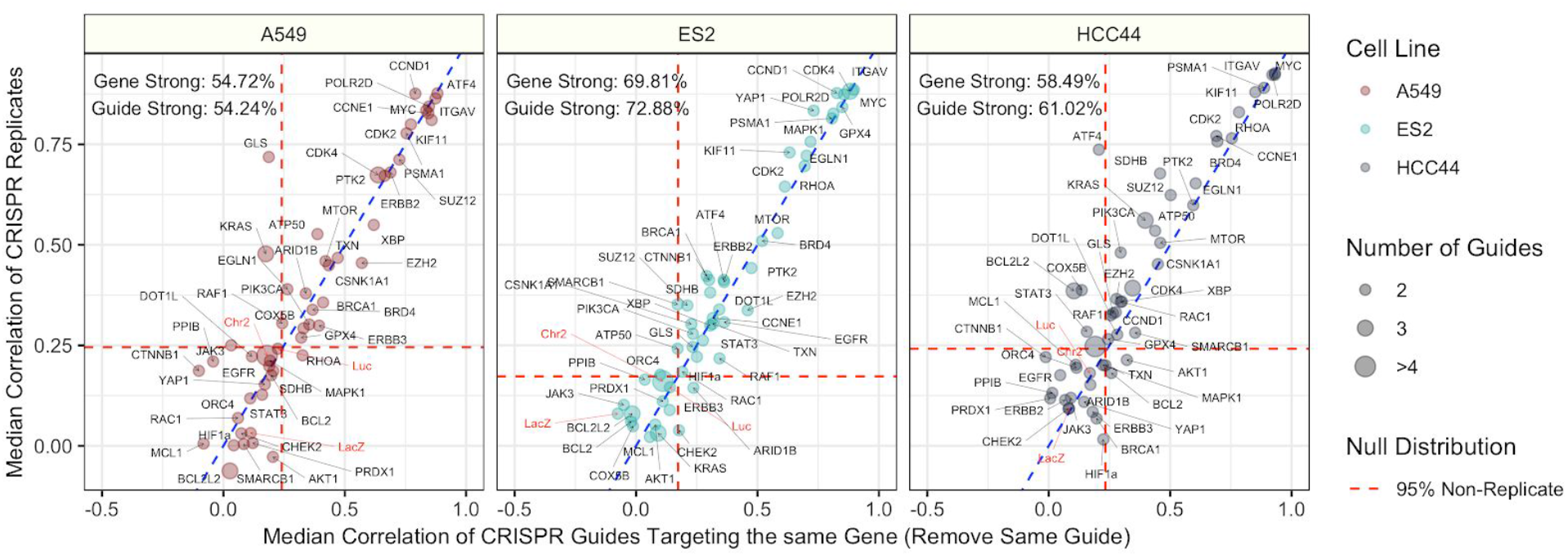
Replicate correlation of CRISPR perturbations. Median pairwise Pearson correlation of CRISPR guide replicate profiles (y axis) compared against Median pairwise Pearson correlation of CRISPR guides targeting the same gene or construct (x axis). We removed biological replicates when calculating the same-gene correlations. The three different cell lines (A549, ES2, and HCC44) are shown in different colors and in different facets of the figure. We generated the profiles by median aggregating CellProfiler measurements for all single cells within each well of a Cell Painting experiment (see Methods for more processing details). The text labels represent the proportion of gene and guide profiles with “strong phenotypes”. In other words, these profiles had replicate correlations greater than 95% of non-replicate pairwise Pearson correlations in the particular cell line. The dotted red line represents this 95% cutoff in the null distribution and the blue dotted line is y = x, which shows a strong consistency across CRISPR guide constructs.

**Supplementary Figure S4.**
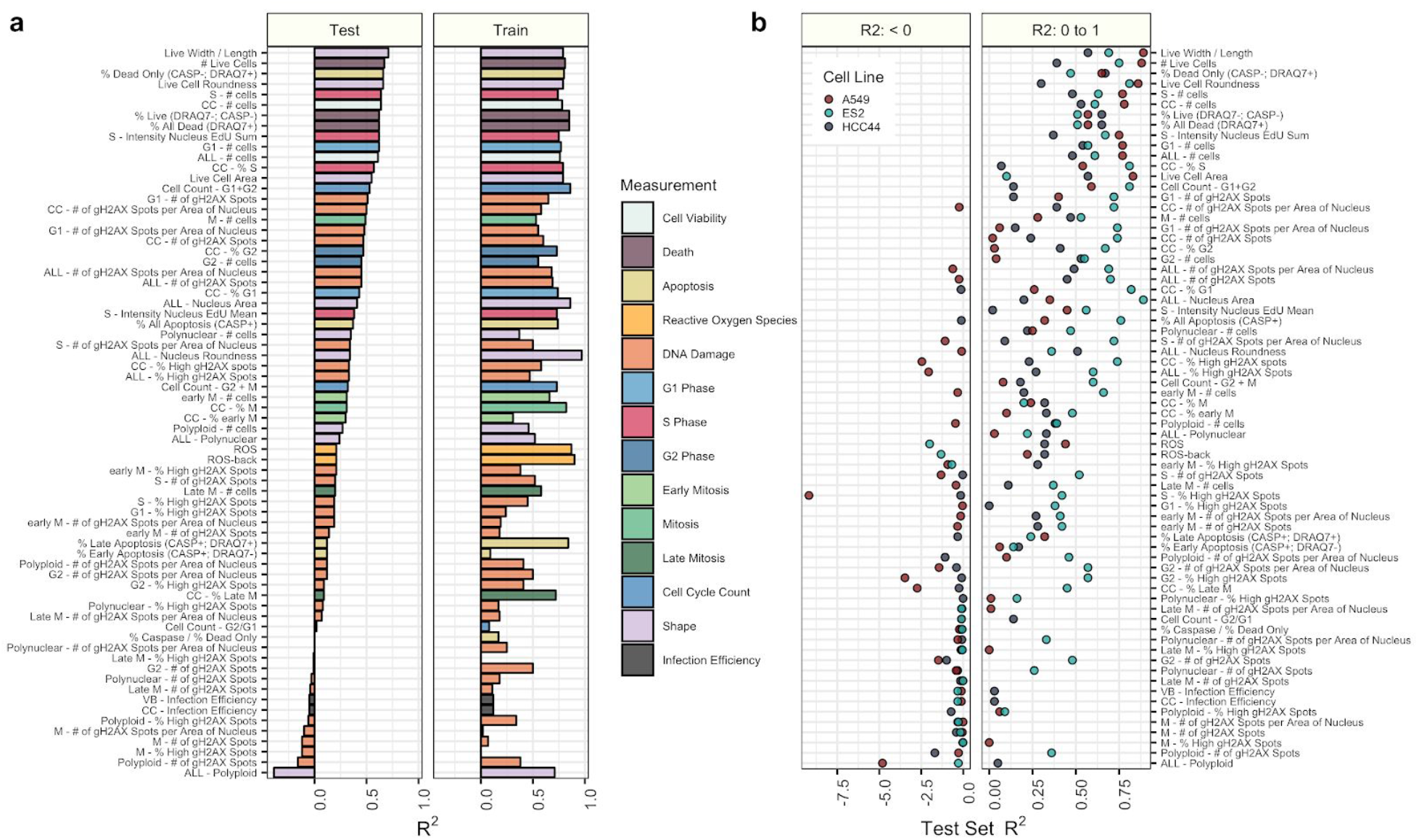
Performance of predicting 70 cell health variables with independent regression models. **(a)** Testing and training performance for each phenotype is shown, sorted by decreasing R^2^ performance on the test set, aggregated across the three cell lines and colored based on the primary measurement metadata (see Supplementary Table S3). **(b)** Test set R^2^ performance for each cell line independently, with the same ordering as in (a). Performance is highly variable across cell lines, with ES2 having the highest performance for most models. Note that the left and right facets of b have different x-axis scales and that the R^2^ values can lie below zero for the test set because the model is learned using the training set.

**Supplementary Figure S5.**
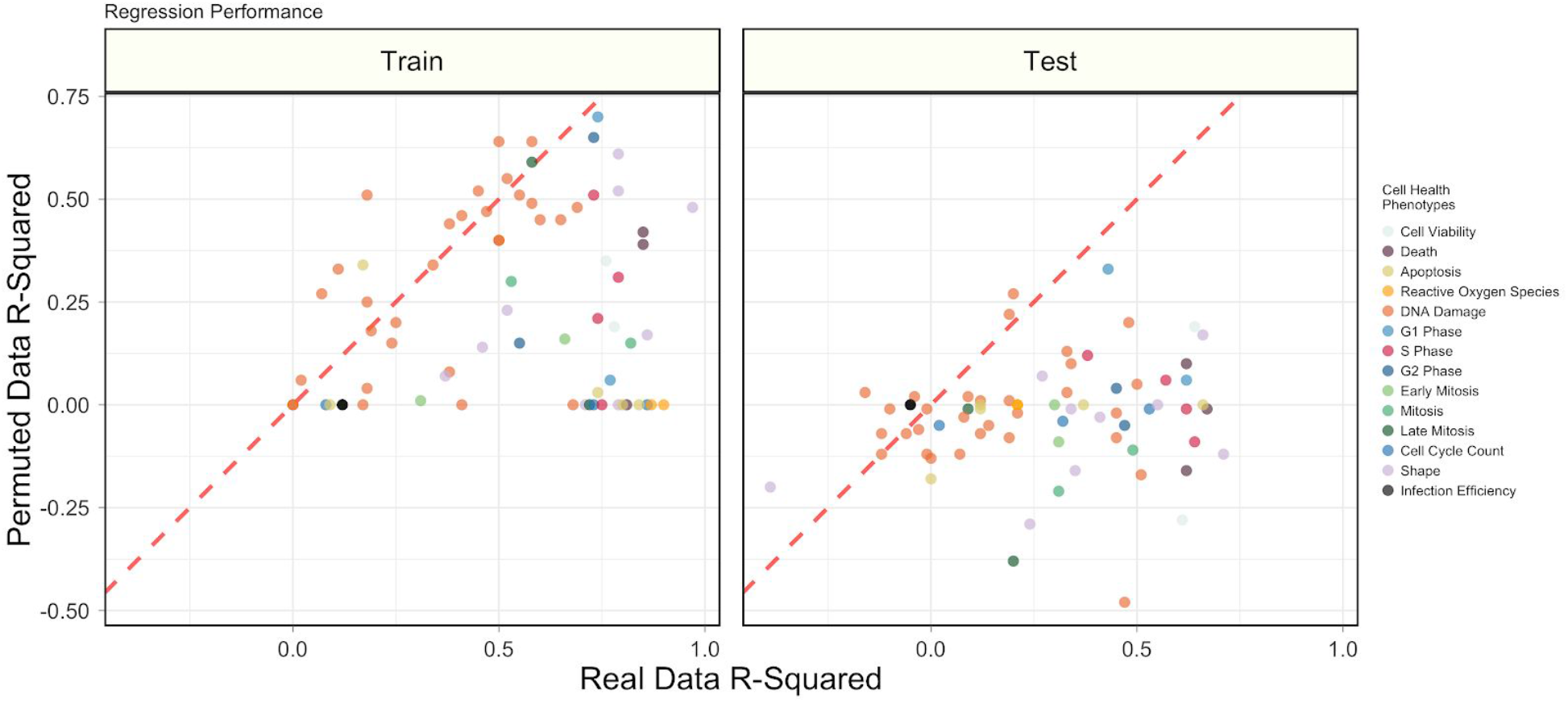
Summarizing model performance compared to permuted data. Performance of all 70 Cell Health assay models in train and test sets using real and permuted data. Data were permuted by randomly shuffling observed morphology features for each column independently (see Methods). Each point represents a Cell Health model, and the color represents the specific phenotype.

**Supplementary Figure S6.**
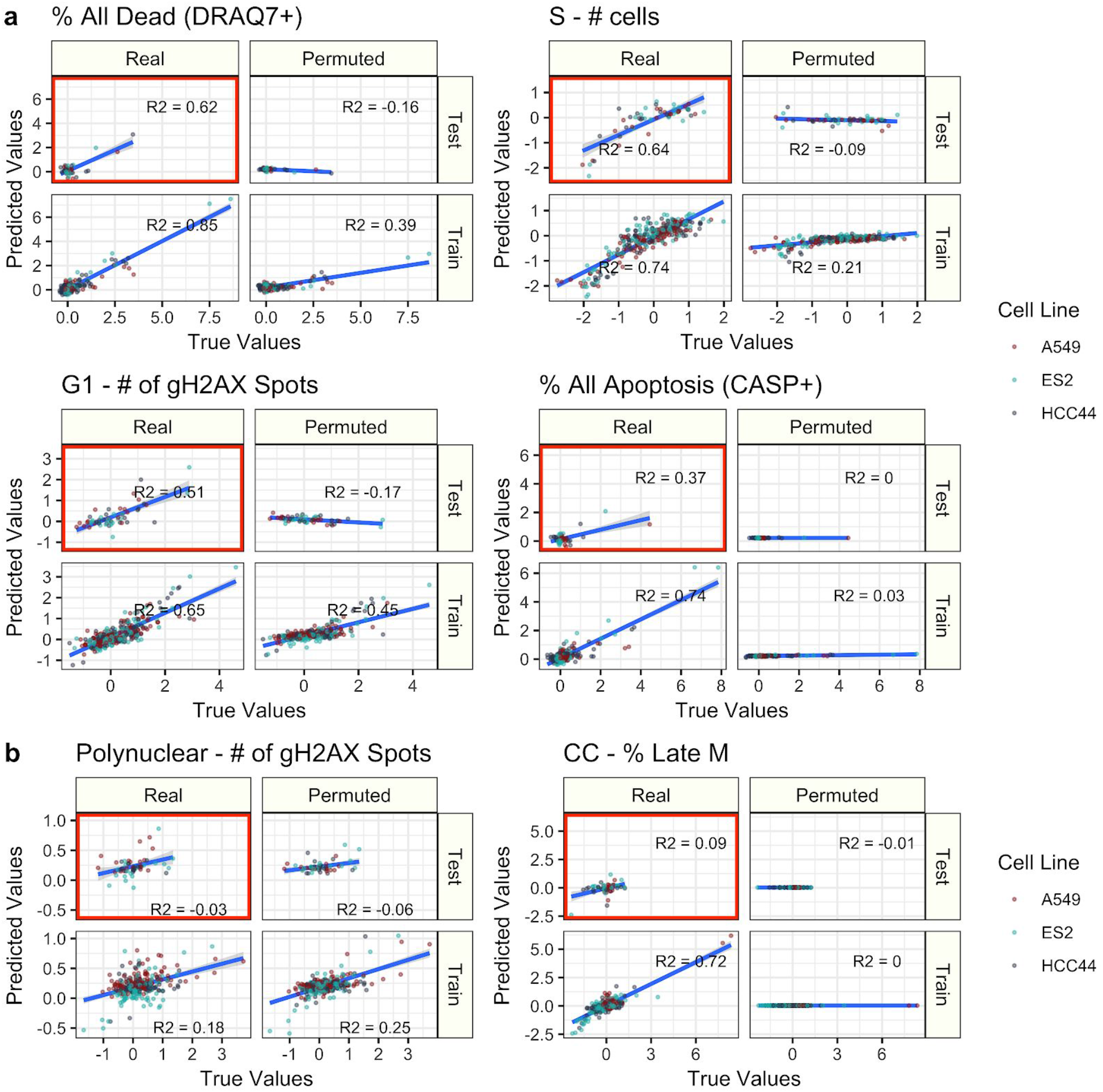
Examples of six models in predicting cell health assay readouts. **(a)** Four high performing models chosen to span different Cell Health assay readouts. **(b)** Predicting two example low performing models. R-squared (R^2^) was calculated using sklearn.metrics.r2_score and is bound by negative infinity to 1. The blue lines represent a smoothed linear model fit and the shading represents a 95% confidence interval.

**Supplementary Figure S7.**
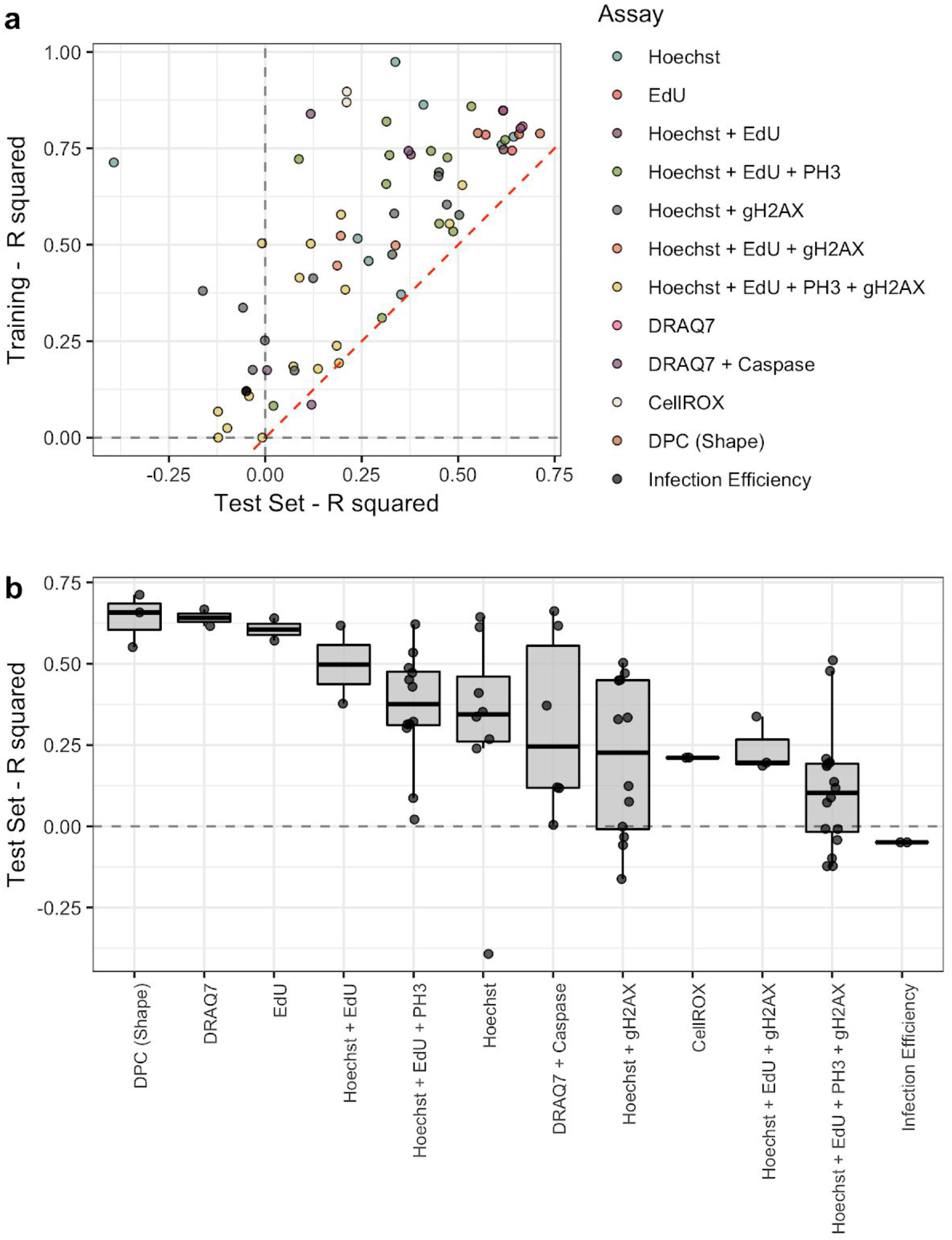
Model performance according to specific cell health reagents. **(a)** Training and testing model performance based on R^2^. The dotted red line is the line y = x and depicts a small amount of model overfitting. **(b)** Test set R^2^ grouped by cell health reagents used to form the cell health indicators. The cell health variables are sorted by median test set R^2^ performance.

**Supplementary Figure S8.**
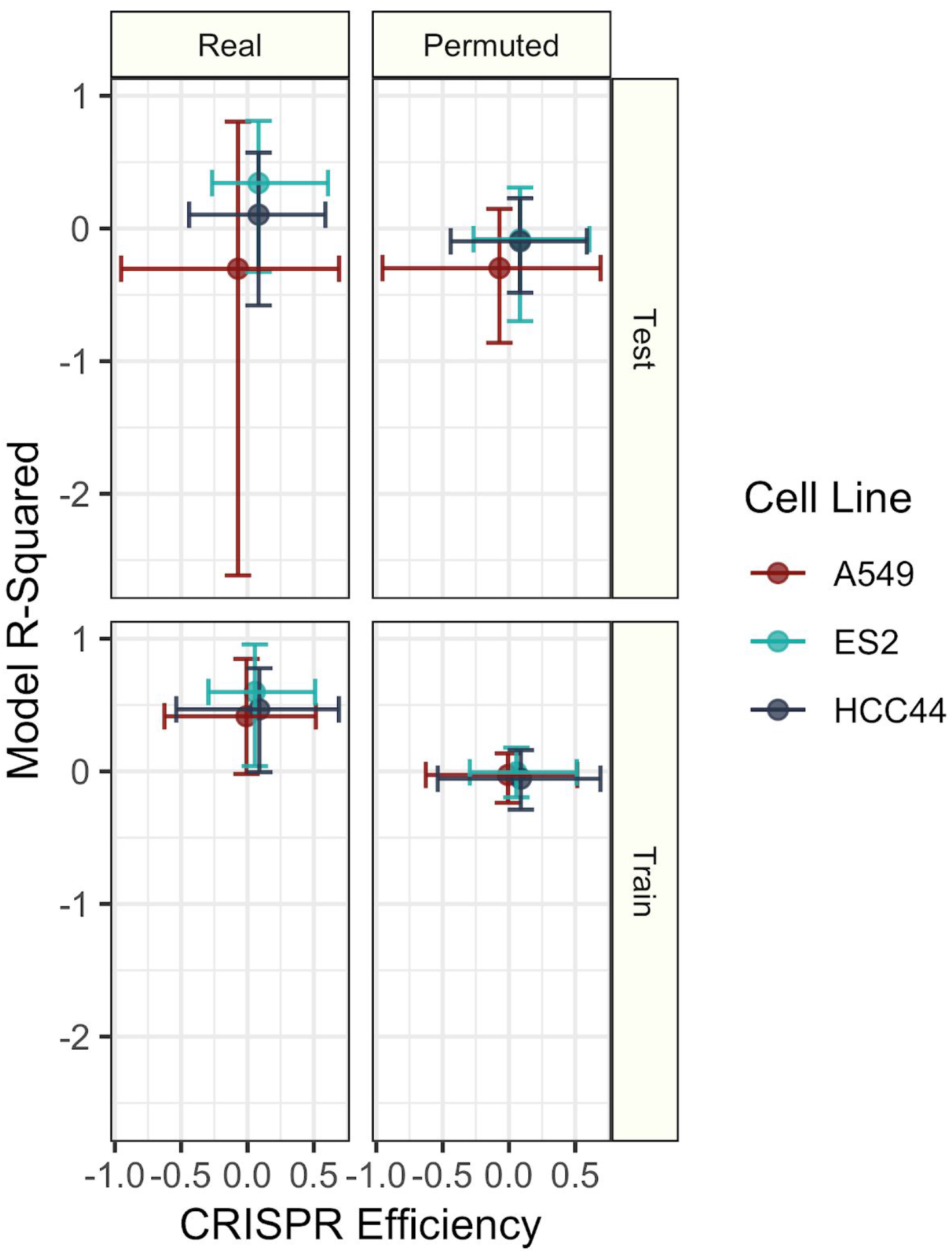
Exploring the relationship between CRISPR infection efficiency and regression model performance. Cell line specific training and testing model R^2^ performance in real and permuted data compared against CRISPR infection efficiency readouts. Infection efficiency is measured by comparing cell count in wells treated with and without puromycin (see Methods). We generated infection efficiency measurements for all individual wells and then aggregated by MODZ to form consensus measurements (see Methods). The points represent mean values and the extended bars represent 5% and 95% of the observed cell-line specific distributions. ES2 has the highest test set model R^2^ and the highest CRISPR efficiency.

**Supplementary Figure S9.**
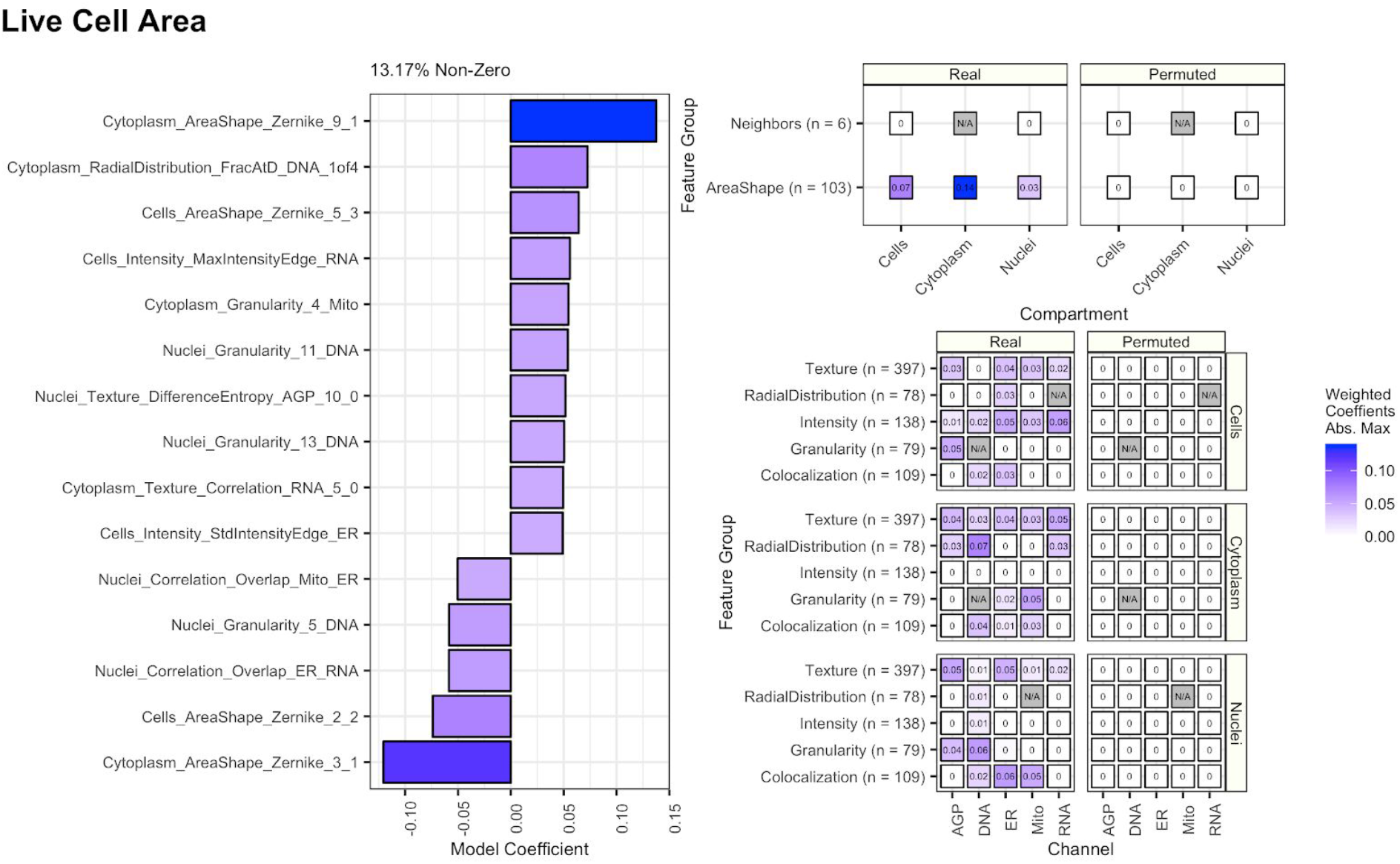
Regression model coefficients for predicting live cell area from digital phase contrast (DPC) measurements. We use the Cell Painting measurements to predict live cell area readouts from DPC measurements as a positive control. As expected, cytoplasm and cell shape contribute to live cell area predictions. **(left)** The top 15 most influential Cell Painting features by absolute value model coefficient. It is important to note that because the machine learning procedure automatically removes many features, not all explanatory features are selected. **(right)** The maximum absolute value model coefficient (weight) for compartments, channels, and feature groups. Coefficients for a model trained with real data is contrasted with a model trained with permuted data. For a complete description of all features, see the handbook: http://cellprofiler-manual.s3.amazonaws.com/CellProfiler-3.0.0/index.html

**Supplementary Figure S10.**
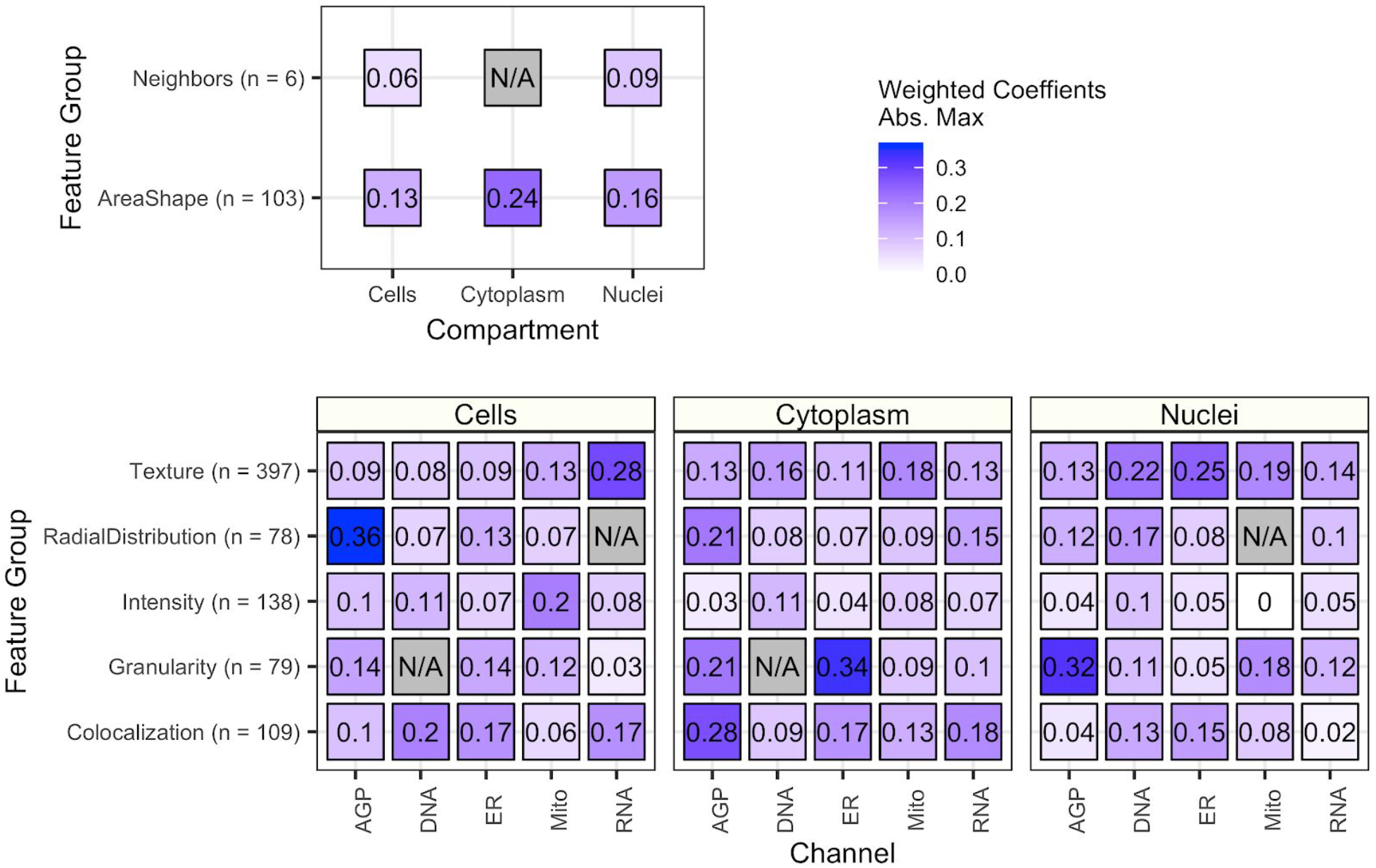
Summarizing Cell Painting feature importance scores for all 70 cell health regression models. Each point represents the maximum absolute value of the model coefficients weighted by test set R^2^ across all Cell Health models. The features are broken down by compartment (Cells, Cytoplasm, and Nuclei), channel (AGP, Nucleus, ER, Mito, Nucleolus/Cyto RNA), and feature group (AreaShape, Neighbors, Channel Colocalization, Texture, Radial Distribution, Intensity, and Granularity). For a complete description of all features, see the handbook: http://cellprofiler-manual.s3.amazonaws.com/CellProfiler-3.0.0/index.html. Dark gray squares indicate “not applicable”, meaning either that there are no features in the class or the features did not survive an initial preprocessing step.

**Supplementary Figure S11.**
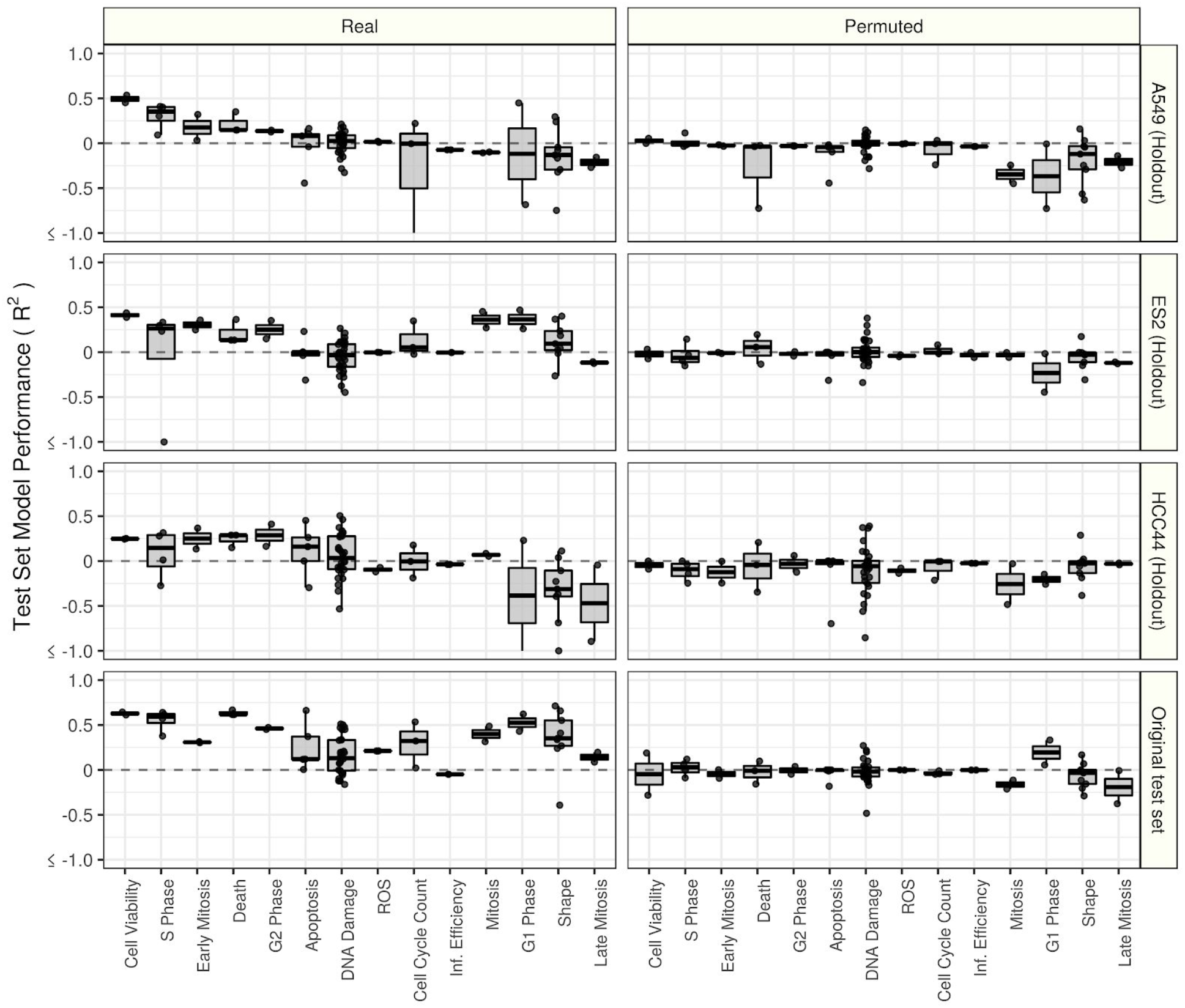
Results from a cell line holdout analysis. We trained and evaluated all 70 cell health models in three different scenarios using each combination of two cell lines to train, and the remaining cell line to evaluate. For example, we trained all 70 models using data from A549 and ES2 and evaluated performance in HCC44. We bin all cell health models into 14 different categories (see Supplementary Table S3 and https://github.com/broadinstitute/cell-health/6.ml-robustness for details about the categories and scores). We also provide the original test set (15% of the data, distributed evenly across all cell types) performance in the last row, as well as results after training with randomly permuted data. This cross-cell-type analysis yields worse performance overall. Nevertheless, despite the models never encountering certain cell lines, and having fewer training data points, many models still have predictive power across cell line contexts. Note that we truncated the y axis to remove extreme outliers far below −1. The raw scores are available on https://github.com/broadinstitute/cell-health.

**Supplementary Figure S12.**
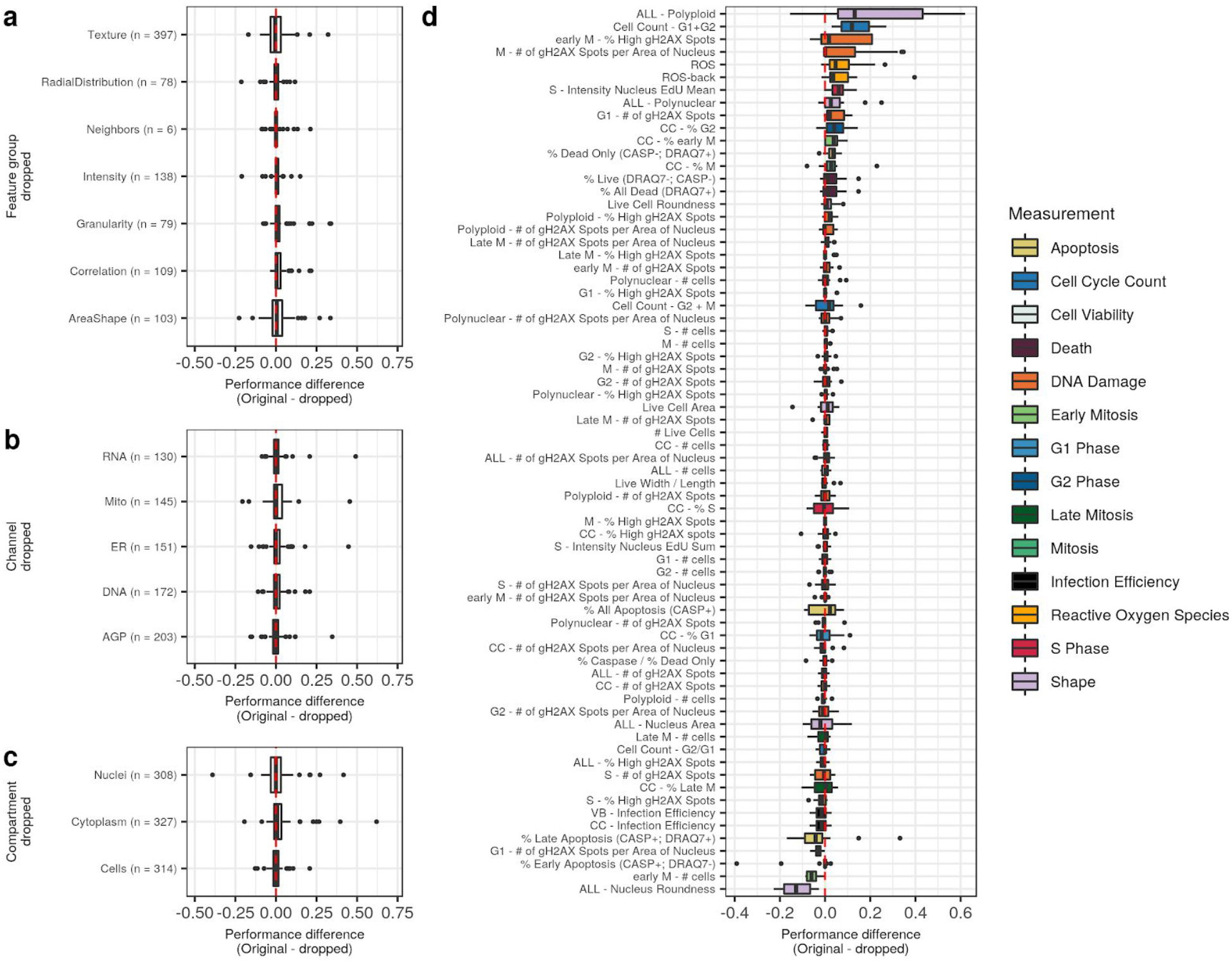
Systematically removing classes of features has little impact on most models’ performance. We retrained all 70 cell health models after dropping features associated with specific **(a)** feature groups, **(b)** channels, and **(c)** compartments. Each dot is one model (predictor), and the performance difference between the original model and the retrained model after dropping features is shown on the x axis. Any positive change indicates that the models got worse after dropping the feature group. **(d)** Individual model differences in performance after dropping features. Each dot is one class of features removed (as in a-c).

**Supplementary Figure S13.**
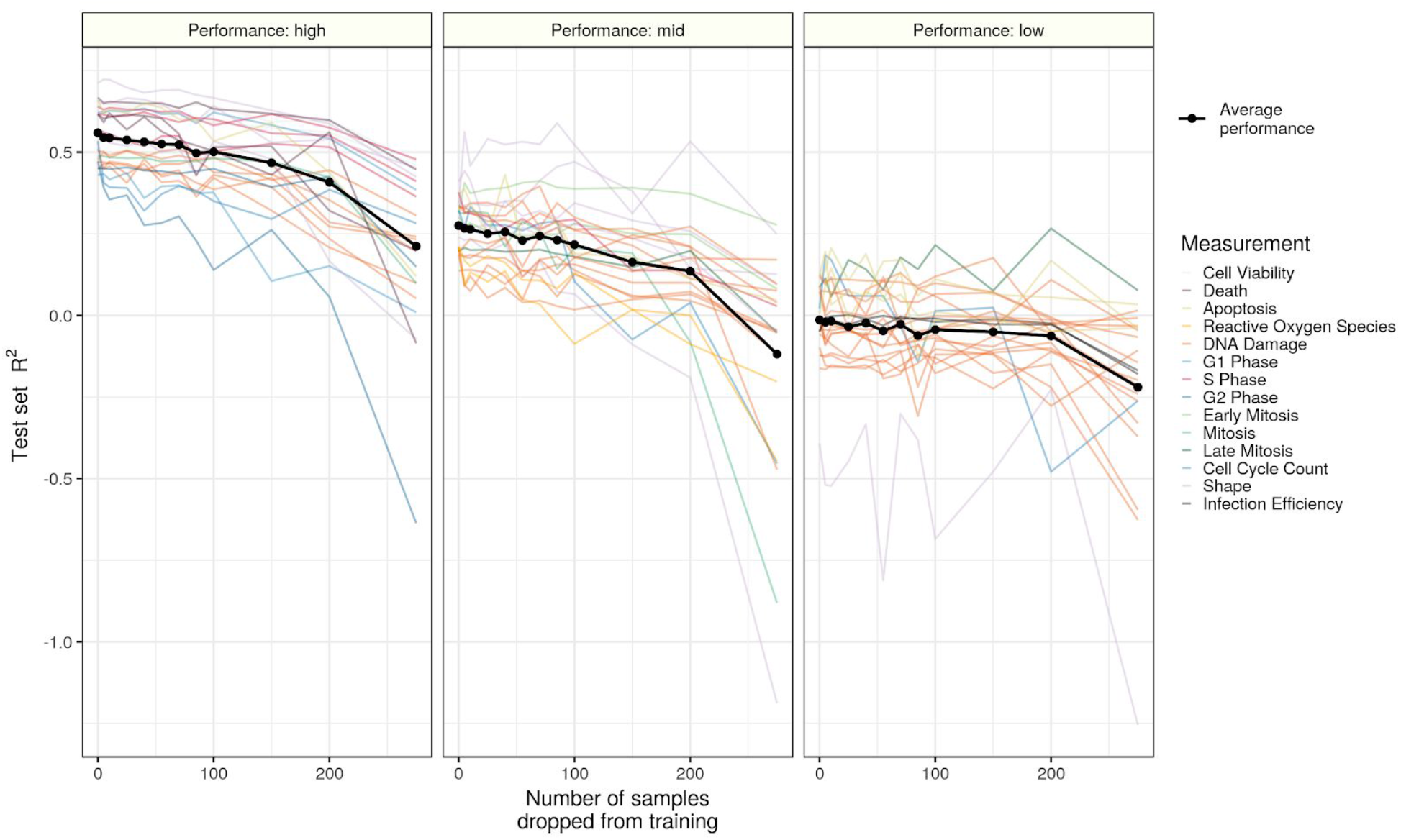
Dropping samples from training reduces test set model performance in high, mid, and low performing models. We determined model performance stratification by taking the top third, mid third, and bottom third of test set performance when using all data. We performed the sample titration analysis with 10 different random seeds and visualized the median test set performance for each model.

**Supplementary Figure S14.**
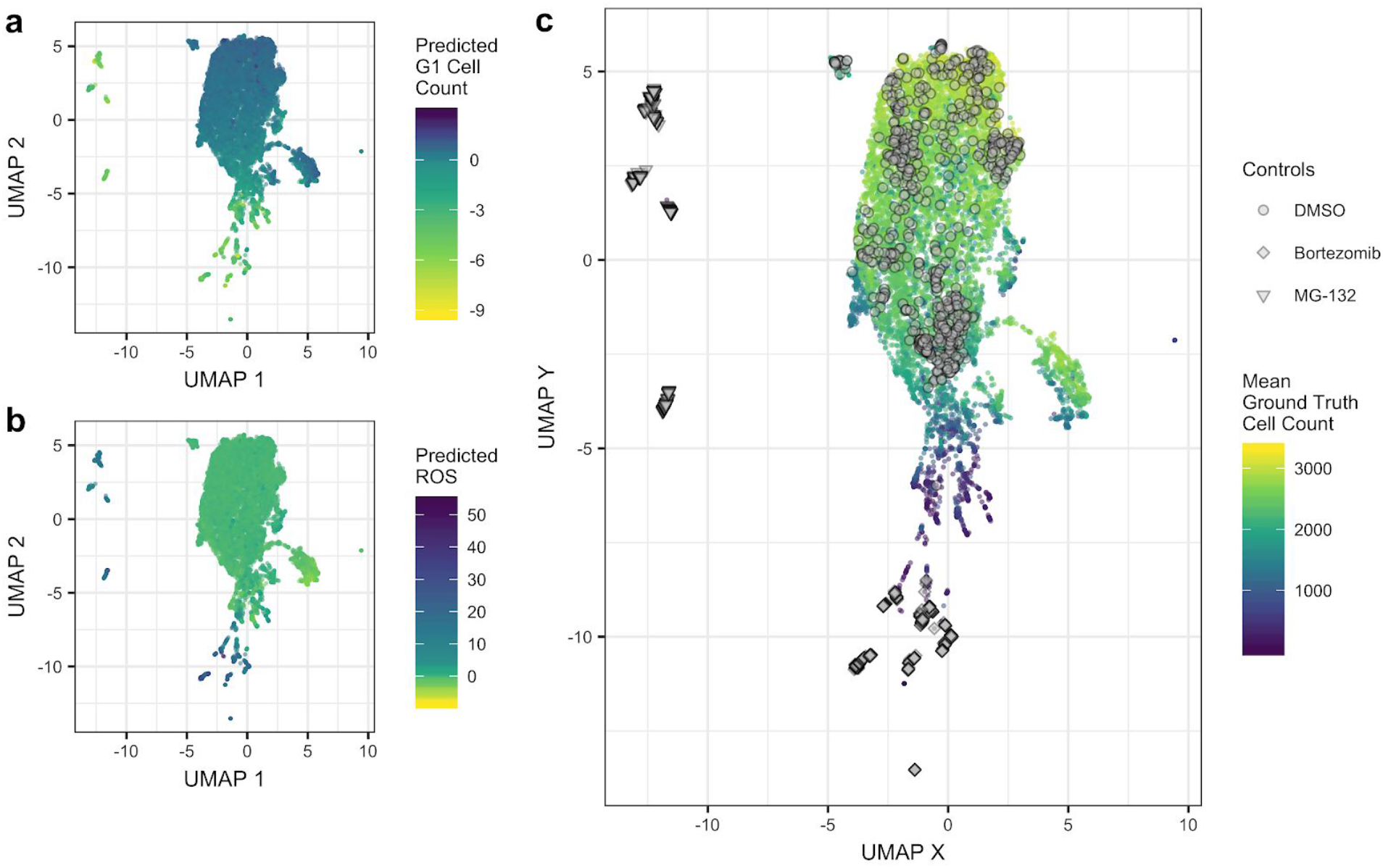
Applying a Uniform Manifold Approximation (UMAP) to Drug Repurposing Hub consensus profiles of 1,571 compounds across six doses. The models were not trained using the Drug Repurposing Hub data. **(a)** The point color represents the output of the Cell Health model trained to predict the number of cells in G1 phase (*G1 cell count*). **(b)** The same UMAP dimensions, but colored by the output of the Cell Health model trained to predict reactive oxygen species (*ROS*). **(c)** In the UMAP space, we highlight DMSO as a negative control, and Bortezomib and MG-132 as two positive controls (proteasome inhibitors) in the Drug Repurposing Hub set. We observe moderate batch effects in the negative control DMSO profiles, based on their spread in this visualization. The color represents the predicted number of live cells. The positive controls were acquired with a very high dose and are expected to result in a very low number of predicted live cells.

**Supplementary Figure S15.**
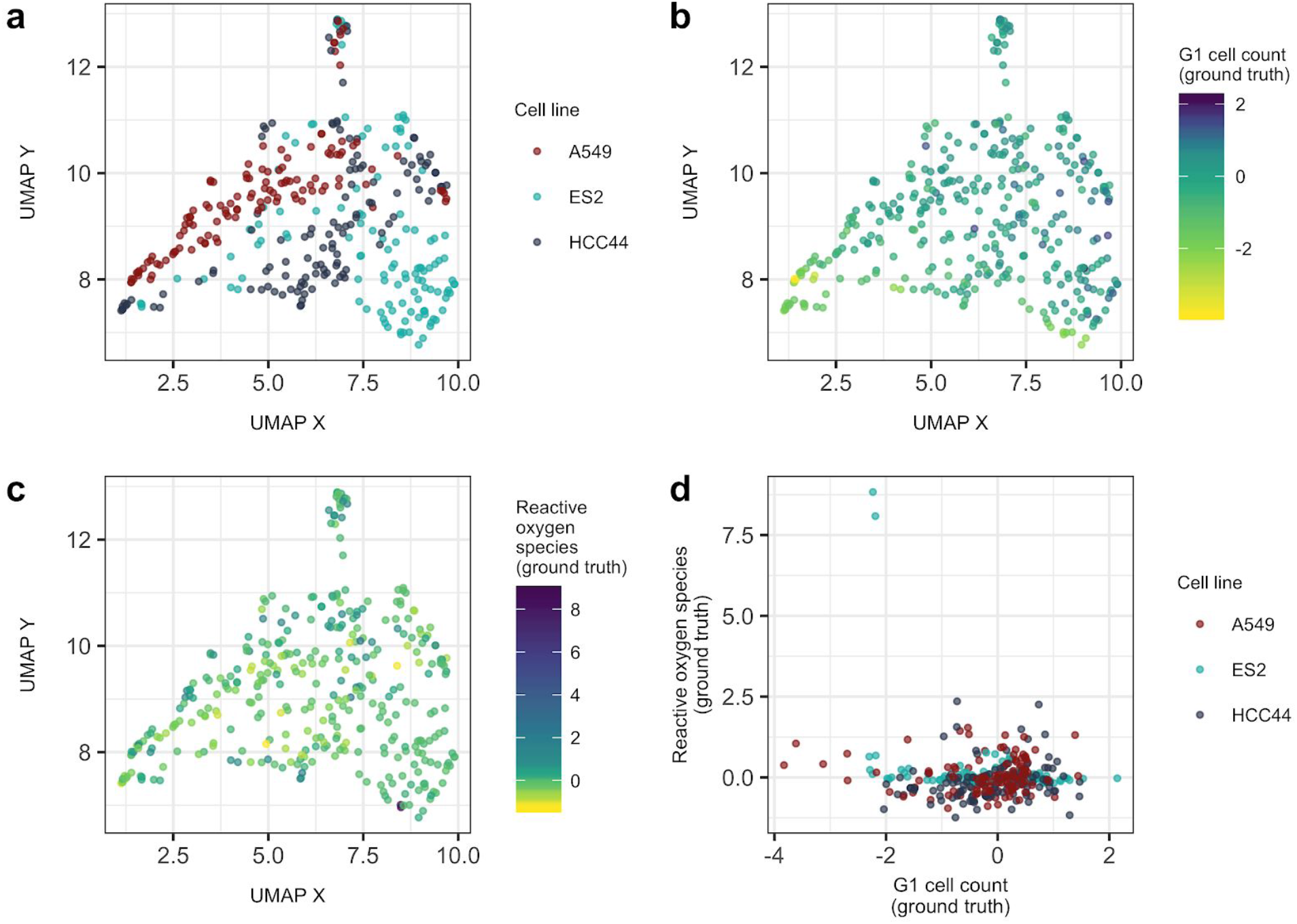
Applying a Uniform Manifold Approximation (UMAP) to the Cell Painting consensus profile data of CRISPR perturbations. UMAP coordinates visualized by **(a)** cell line, **(b)** ground truth G1 cell counts, and **(c)** ground truth ROS counts. (d) Visualizing the distribution of ground truth ROS compared against G1 cell count. The two outlier ES2 profiles are CRISPR knockdowns of GPX4, which is known to cause high ROS.

